# Integrating linguistics, social structure, and geography to model genetic diversity within India

**DOI:** 10.1101/164640

**Authors:** Aritra Bose, Daniel E. Platt, Laxmi Parida, Petros Drineas, Peristera Paschou

## Abstract

India represents an intricate tapestry of population substructure shaped by geography, language, culture and social stratification. While geography closely correlates with genetic structure in other parts of the world, the strict endogamy imposed by the Indian caste system and the large number of spoken languages add further levels of complexity to understand Indian population structure. To date, no study has attempted to model and evaluate how these factors have interacted to shape the patterns of genetic diversity within India. We merged all publicly available data from the Indian subcontinent into a data set of 891 individuals from 90 well-defined groups. Bringing together geography, genetics and demographic factors, we developed COGG (Correlation Optimization of Genetics and Geodemographics) to build a model that explains the observed population genetic substructure. We show that shared language along with social structure have been the most powerful forces in creating paths of gene flow in the subcontinent. Furthermore, we discover the ethnic groups that best capture the diverse genetic substructure highlighted by COGG. Integrating data from India with a data set of additional 1,323 individuals from 50 populations we find that Europeans show shared genetic drift with the Indo-European and Dravidian speakers of India, whereas the East Asians have the maximum shared genetic drift with Tibeto-Burman speaking tribal groups.

## Introduction

The genetic structure of human populations reflects gene flow around and through geographic, linguistic, cultural, and social barriers [16], [59]. The intricate tapestry of population substructure and complexity in India undoubtedly showcases the interplay among them; 3,200 km from North to South, complex topography with elements ranging from the Himalayas to the Thar desert, plateaus and rain forests, almost 800 spoken languages, a long history of migrations and invasions and a strict caste system imposing endogamy are factors that have shaped extant human genetic diversity within India.

The strata within India can be summarized into the so-called backward castes and forward castes ([24]), while 8.2% of the total population belongs to tribes (1991 census) representing minorities that are unassimilated into the caste system. The tribes in India continue to live in forest hills and naturally isolated regions with a largely hunting-gathering subsistence mode. They practice endogamy, governing mate-exchange within local groups [64]. On the other hand, the caste system is a rigorous social hierarchy of endogamous groups in which individuals are born into [43], [68]. Prior to the establishment of the caste system there was wide admixture among them, which came to an abrupt end 1,900 to 4,200 years before present [39]. Historically, the so-called forward castes have been associated with socio-economic privileges while the backward castes and tribal groups faced social segregation [24]. Although discrimination on the basis of caste was abolished by the Indian constitution in 1950, this strict social structure has existed for thousands of years [63].

Numerous studies have attempted to dissect the genetic components and origins of Indian populations [7], [34], [55], [8], [14], [53], [37], [4], [39], [9], [58], [46] along with ancient individuals from Central and South Asia [40]. Studies of Indian populations based on groupings of tribal versus non-tribal, geographic regions, or linguistic affiliation have shown that the observed genetic structure resulted from admixture of five ancestral populations. These are Ancestral North Indians, which loosely captures Indo-European (IE) speakers in Northern India; Ancestral South Indians, who are mostly Dravidian (DR) speakers of Southern India; Ancestral Austroasiatic with Austroasiatic (AA) speakers of Central and Eastern India; Ancestral Tibeto-Burman speakers constituted of Tibeto-Burman (TB) speakers in Northeast and the tribal populations from Andaman archipelago (AND) [9]. Although, the Great Andamanese are language isolates, typologically and geneaologically different from other Andamanese populations, Jarawa and Onge and their spoken language is considered as the sixth language family of India [1]. However, to date, no study has attempted to model how different spatio-cultural features acted in concert in order to create the observed genetic structure across the Indian subcontinent and to evaluate the relative contribution of each factor.

Earlier attempts to investigate the covariance of allele frequencies and non-genetic factors on genetic structure, either depended heavily on assumptions and a computationally expensive Bayesian framework [13] or did not provide any statistical significance or feature selection to identify the most relevant structure-related factors [57]. To dissect the population substructure in Indian populations, we designed a quantitative framework for the evaluation of the relative contribution of geodemographic features such as geography, spoken language and social structure to the architecture of the genetic pool of human populations. Our work provides a general model that may be used to study the significance of each underlying factor on the genetic substructure of a given population.

## Materials and Methods

### Study design and datasets

We used PLINK1.9 ([17]) to assemble genome-wide data for 891 samples from 90 well-defined sociolinguistic groups (**Figure 1**; **Supplementary Table S1**) genotyped on 47,283 autosomal SNPs. These samples were collected from various sources ([53]; [37]; [19]; [39]; [9]) with the consent of the corresponding authors. We created subsets of this dataset in order to construct an equal representation of social groups, language families and geographical locations for this study and tested for correlation between genetics and geography along with sociolinguistic features. The normalized subset (See **Supplementary Notes** for details) for which we have reported results on COGG, contains 368 samples from 33 populations genotyped on 47,283 SNPs (**Supplementary Table S1B**). We converted all data to the same build (hg19) using LiftOver from the UCSC Genome Browser ([30]) before merging the data. Further quality control such as filtering out variants with missing call rates exceeding 5% and minor allele frequency (MAF) of at least 5% was performed in PLINK ([50]; [17]).

**Figure 1.**
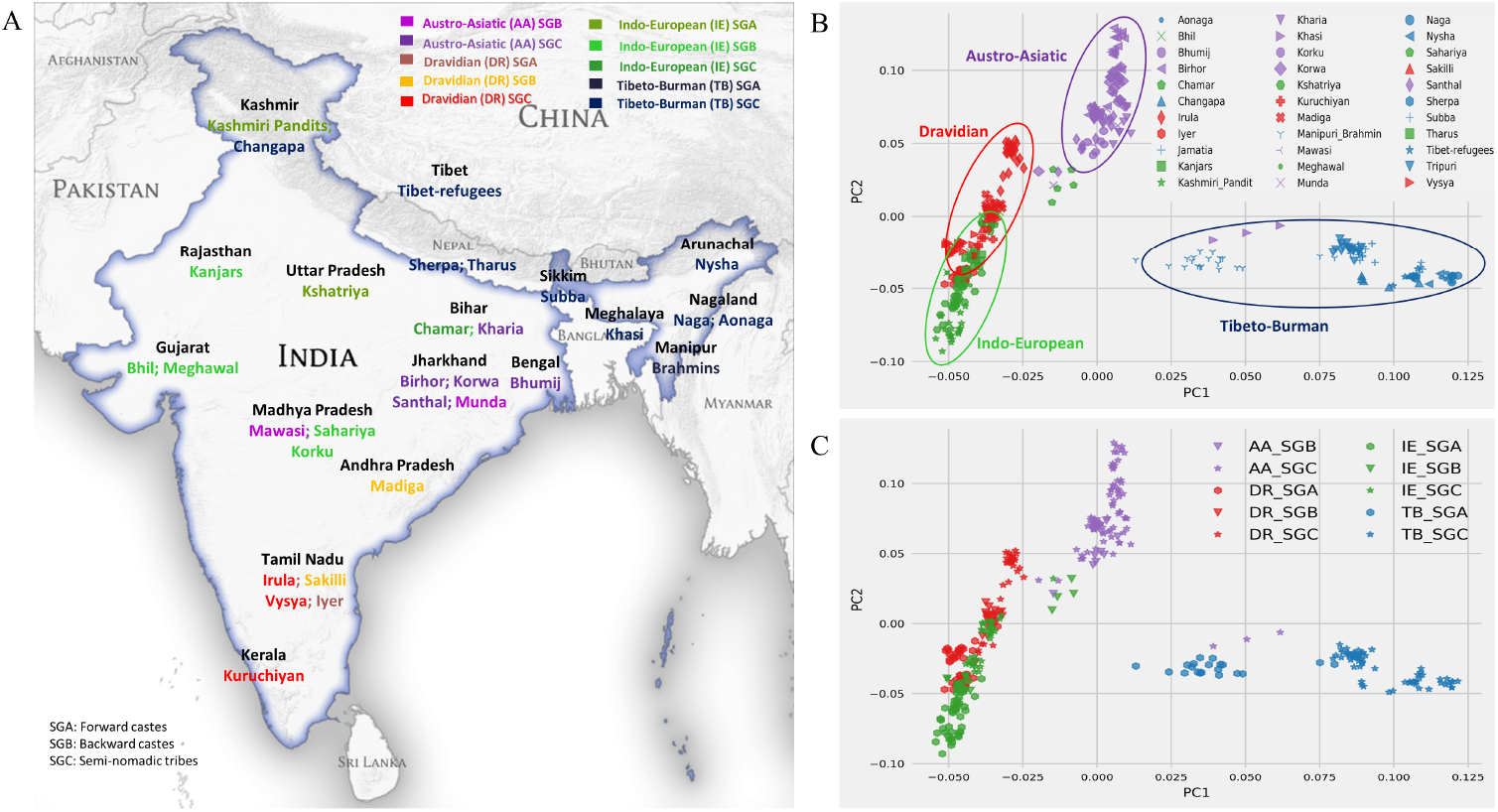
**A**. Map of India showing the pan-Indian locations of the 368 individuals in the normalized subset across 33 well-defined population groups, 47,283 SNPs, (see Supplementary Figure 1A for the entire data set of 90 ethnic groups across India). The populations are colored by their sociolinguistic group. **B**. Top two PCs of the normalized data set show clustering by language groups. **C**. PCA plot colored and marked by sociolinguistic groups affirming the stratified gene flow in Indian subcontinent.

We merged 1,323 individuals across 50 populations from Eurasia and Southeast Asia, collected from various publicly available sources such as HGDP ([15]), the Estonian Biocenter ([10]; [69]; [25]; [27]; [32]; [51]; [70]) and the Allele Frequency Database (ALFRED) ([52]) (**Supplementary Table S1C**) with our normalized Indian dataset to create a merged data set of 1,691 samples from 83 population groups genotyped on 42,975 SNPs overlapping between all data sets (**Supplementary Table 1**).

### Mantel Tests

We ran Mantel tests on the normalized Indian data set by computing pairwise *F*_*ST*_ distances between 33 groups using PLINK 1.9 ([17]). We performed 10,000 permutations and estimated Spearman’s correlation, acknowledging the caveat of overestimation of p-values obtained from the tests ([28]).

### PCA and LDA

We used TeraPCA ([12]) to perform PCA on our datasets after pruning for LD structure by setting `--indep-pairwise` 50 10 0.4 in PLINK 1.9 ([17]). We checked for outliers (using EIGENSTRAT’s ([49]) outlier detection method) in the PCA plot (**Supplementary Figure 2A**) and removed three outliers, each one from TB speakers Jamatia, Tripuri and Sherpa.

We implemented Rao’s Discriminant Analysis which is directly based on Fisher’s Linear Discriminant Analysis (**Supplementary Note**).

### COGG and feature selection using Orthogonal Matching Pursuit

COGG stands for Correlation Optimization of Genetics and Geodemographics and maximizes the correlation between one of the top two principal components (for more PCs see CCA section in **Supplementary Note**) and the Geodemographic matrix, aimed to model genetic structure within India, consists of nine features (columns) corresponding to geographical coordinates, social groups and language information encoded as indicator variables. We restrict our encoding into three social groups (A, B and C) as described before. **u** is the vector containing either one of the top two principal components, computed by TeraPCA ([12]); the Geodemographic matrix is mean-centered and denoted as **G** ∈ ℝ^*m*×*k*^, including *k* features across *m* individuals. The social group (A, B, C) and language (AA, DR, IE, TB) encoding was performed as 1 if the sample belongs to that social group (or language) and 0, otherwise.

Let **a** be the *k*-dimensional vector whose elements are *a*_1_ … *a*_*k*_ (in our case, *k* = 9). COGG solves the following optimization problem (see **Supplementary Note** for details):

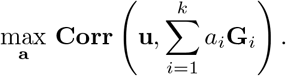

Recall that **G**_*i*_ denotes the *i*-th column of **G** as a column vector. Let 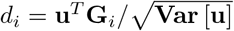 for *i* = 1 … *k* and let **d** be the vector of the *d*_*i*_’s. Also, let 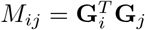 for all *i, j* = 1 … *k* and let **M** be the matrix of *M*_*ij*_. Then the optimizer for COGG is given by

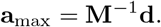

We also check for statistical significance of the maximum squared Pearson correlation coefficient *r*^2^, returned by COGG, by randomly permuting the columns corresponding to social factors and languages in **G** in 1,000 iterations and calculating **a**_max_ for each iteration; we report the histogram of the resulting *r*^2^ values (**Supplementary Figure 6**).

We used a greedy feature selection algorithm described in ([41]) to select features of the Geodemographic matrix **G**. We obtain two sets, *S*_1_ and *S*_2_ of the three most significant quantitative features from the nine features in **G**, one for PC1 and the other for PC2. The algorithm is described in detail in the **Supplementary Note**. In short, it selects the column which results in the maximum *r*^2^ value from **G** and then projects **G** (and **u**) on the subspace perpendicular to the selected column in order to form **G**′ (and **u**′). We iterate the process until we remove the required number of features from **G**.

All the values returned by this method are statistically significant, as random permutations of the elements of the two feature sets return negligible *r*^2^. We considered all 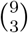 possible sets of three features exhaustively and concluded that, out of all possible sets, using only *S*_1_ and *S*_2_ would return the maximum correlation to PC1 and PC2 respectively.

### Ridge Leverage Scores

We devised a simple method based on the Ridge Leverage Score (RLS) statistic in order to identify Indian populations that maximally contribute to the genetic diversity within the Indian sub-continent. We considered the genotype data, denoted by mean-centered (by SNPs) matrix **Z** ∈ ℝ^*m*×*n*^ where *m* is the number of individuals and *n* is the number of markers in the pan-Indian data set of 90 Indian populations (891 individuals) and 47,283 SNPs. Since we are interested in the median RLS statistic as the representative of a population, including groups of larger sample size would not introduce any biases, so there was no need for normalization. We also considered the mean-centered Geodemographic matrix **G**. Our analysis procedure based on the RLS statistic has four steps:

- We apply the RLS algorithm separately to the matrices **Z** and **G** to find their corresponding row ridge leverage scores, denoted by 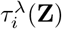 and 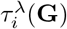, respectively, for *i* = 1 … *m*.
- We grouped the RLSs by population groups to obtain a single score (median RLS) per group. If there are *T* = {*t*_1_, *t*_2_, …, *t*_*T*_} populations in the entire set of the Indian populations (|*T*| = 90 in this case), then we obtain |*T*| RLSs in this manner, one per population *t*_*i*_, defined as the |*T*| × 1 vectors 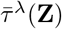 and 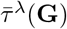.
- Next, we compute an additive RLS for each population after normalizing the vectors obtained in the last step. This additive RLS highlights the significant rows (in our case, Indian populations), across both the genotype and the Geodemographic matrices. We define this consolidated additive RLS as,

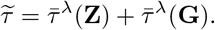
- Finally, we sort the entries of 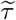 in descending order to obtain a set of representative populations.

### Estimating population admixture and meta-analysis

We used the ADMIXTURE v1.22 software ([3]) for all admixture analyses. Prior to running ADMIXTURE, we pruned for LD using PLINK ([17]) by setting `--indep-pairwise` 50 10 0.8. To determine the optimal number of ancestral populations (*K*), we varied *K* between two and eight performing iterations until convergence for each value of *K* (performing this procedure eight times).

We also performed a quantitative analysis (**Supplementary Note**) of ADMIXTURE’s output using a method described and implemented in ([60]). To compute the shared ancestry between populations **X** and **Y**, we create two matrices **P**_**X**_ ∈ ℝ^*x*×*K*^ and **P**_**Y**_ ∈ ℝ^*y*×*K*^ containing the estimates from ADMIXTURE, where *x* and *y* are the numbers of samples in **X** and **Y** respectively. Thereafter, we project **P**_**X**_ onto the subspace spanned by **P**_**Y**_. In other words, we take the top *p* eigenvectors of **P**_**X**_, **V**_**X**_ and perform the following to find the shared ancestry between **X** and **Y**,

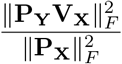

We designed a color-coding scheme to better visualize the results, where the highest shared ancestry between two populations is black and the lowest shared ancestry is white. All intermediate values of shared ancestry follow a gradient from white to black.

### Three population statistics and network analysis

We used ADMIXTOOLS ([47]) to compute *f*_3_ statistics and used YRI (Yorubans) as an outgroup population. We set the significance thresholds for z-score as |*Z*| > 3.

To better visualize and understand the connection between the populations included in our study, we performed a network analysis on the meta-analysis results of ADMIXTURE, using a method presented by a previous study ([44]). We varied the nearest neighbors (NN) of the graph (**Figure 2**) from four to eight for NN = 5. As we can have 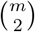 number of edges for an undirected graph with *m* nodes, we allow edges to the graph (**Figure 2**) until all the *n* populations (nodes) appear in the graph with their corresponding NNs sorted by decreasing edge weight (shared ancestry). For Figure 2 we obtained the top 40% of all edges using this criterion.

**Figure 2.**
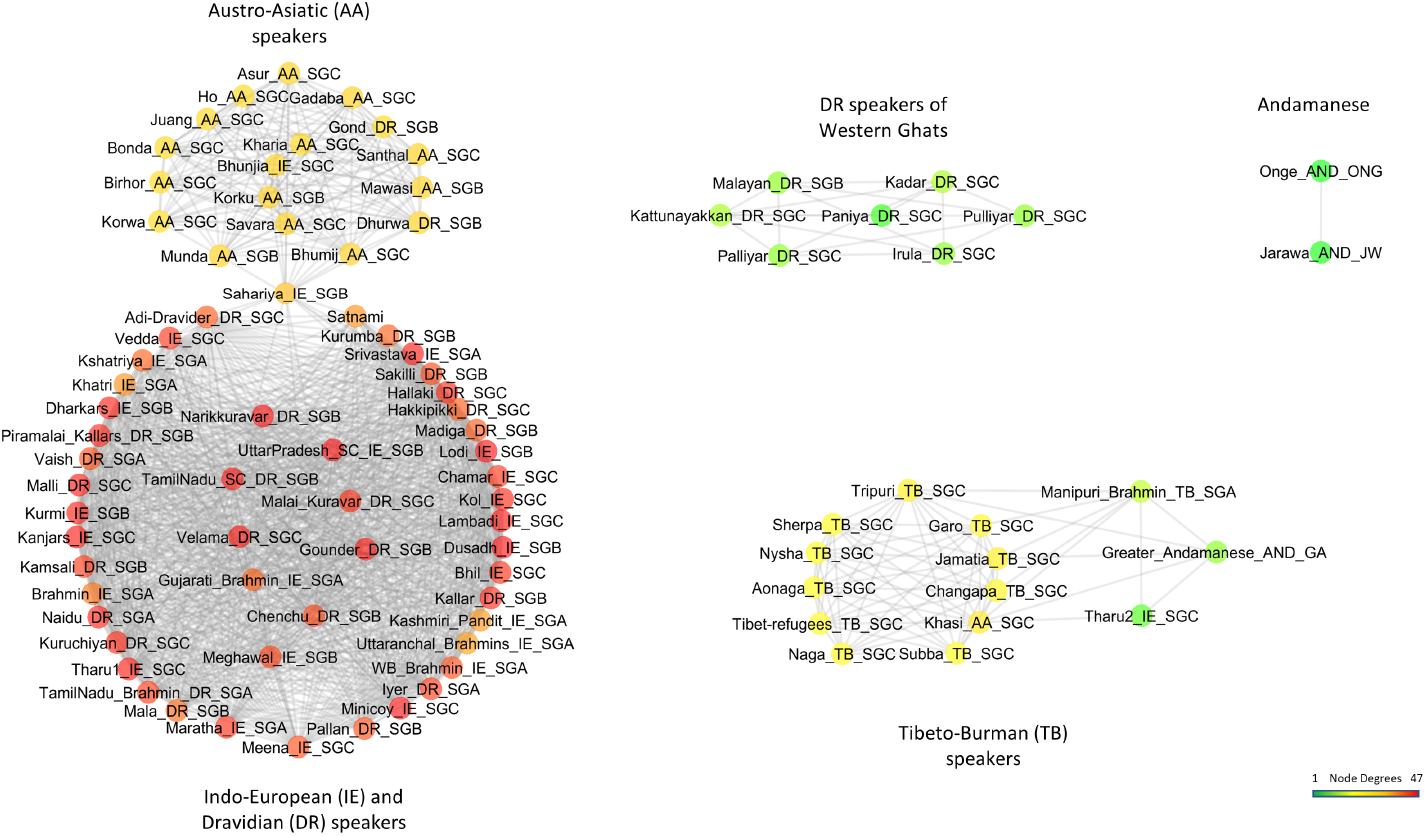
Network of 90 Indian populations (891 individuals) based on shared ancestry as defined by meta-analysis of ADMIXTURE results. Only the top 40% of edges (most related) populations are shown here (see Methods for details).

**Figure 3.**
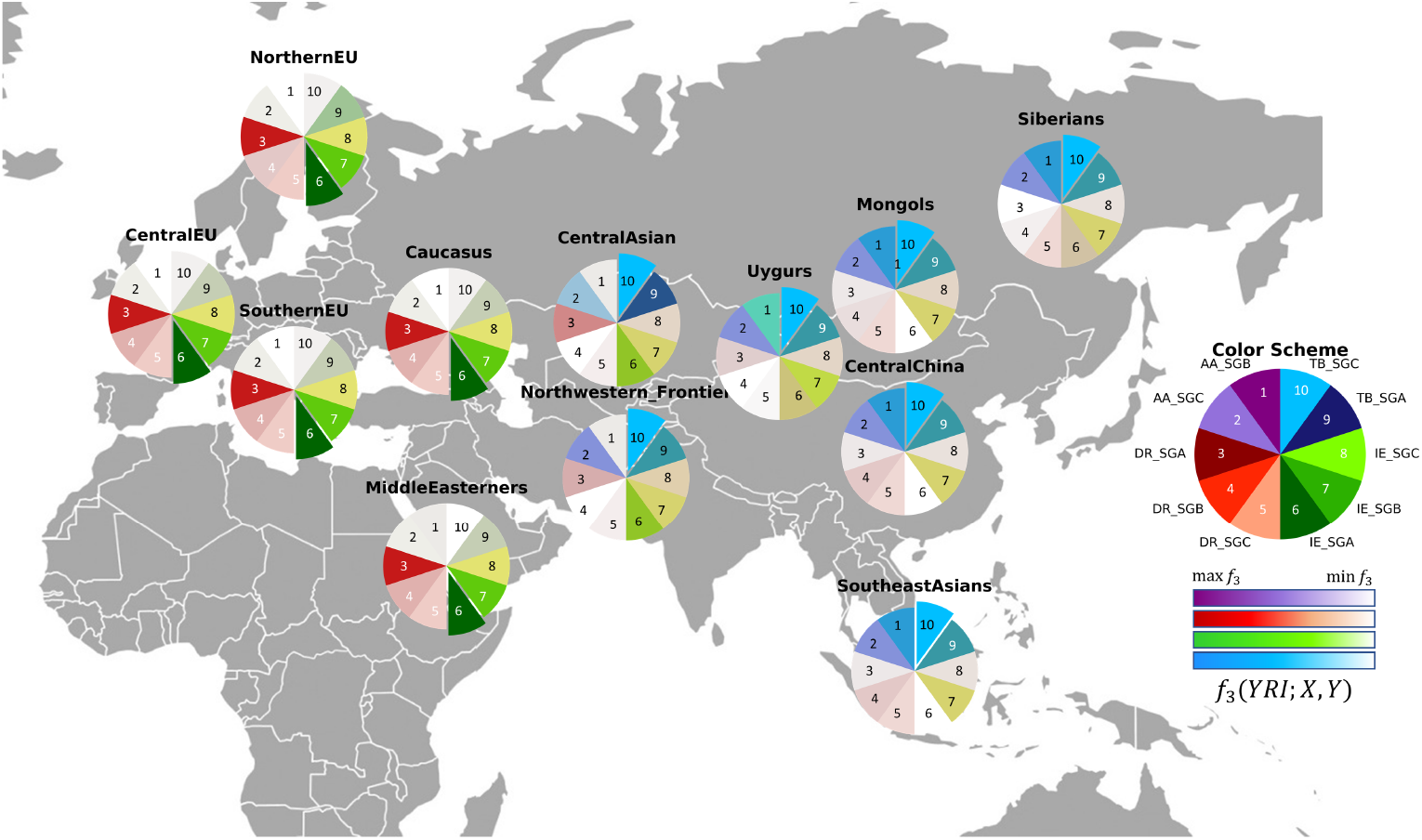
Shared genetic drift between 33 Indian populations (denoted by X) and 50 Eurasian/East Asian populations (denoted by Y) as estimated by *f*_3_ statistics with Yoruba as an outgroup *f*_3_(YRI;X,Y). Europeans show shared genetic drift with the IE and DR speakers of India, whereas the East Asians have the maximum shared genetic drift with TB speakers from SGC (tribal groups). The darkest colors correspond to greatest portions of shared genetic drift with Indian populations. Full results can be found in Supplementary Table S4.

## Results and Discussion

### Description of Compiled Data Sets

We begin by briefly introducing the different data sets that are presented throughout our analysis (**Supplementary Table 1**). We initially compiled a pan-Indian data set of 891 individuals across 90 population groups (**Supplementary Table 1A**) and 47,283 SNPs from various sources [53], [37], [19], [39], [9]. This data set presented unequal representations of the five language families IE, DR, AA, TB and AND as well as uneven distribution across social groups and geographical regions. To create a normalized subset across these spatio-cultural features we selected a subset of 33 populations spanning 368 individuals (**Supplementary Table 1B**) from the pan-Indian data set and used it for the COGG and subsequent feature selection analyses. For other analyses such as the RLS statistic identifying representative ethnic groups contributing to the genetic diversity in India and relationship between sociolinguistic groups, we used the pan-Indian data set. Furthermore, in order to interrogate the shared ancestry between Eurasia and the Indian sociolinguistic groups, we merged the normalized subset with 1,323 individuals from 50 populations and 42,975 SNPs across Eurasia (**Supplementary Table 1C**). For the outgroup f3 analysis we present later in this section, we also used 124 samples of Yorubans in Nigeria (YRI) in 1000 Genomes data set [5] and merged it with the Eurasian data set.

### Geography versus population structure within India

Studies of populations in different parts of the world, have previously shown that, when top PCs are extracted from genome-wide genotypes, individuals from the same geographic region cluster together with the PCs being well correlated with geographic coordinates, namely longitude and latitude [33], [54], [22], [45]. For instance, Novembre et al. (2008) [42] showed that within Europe, the squared Pearson-correlation coefficient *r*^2^ between the top singular vector of the genetic covariance matrix vs. latitude (North-South) is equal to 0.77 and 0.78 for the second singular vector of the same matrix vs. longitude (East-West). In order to explore whether Indian genetic information mirrors geography, we computed the top two PCs using TeraPCA [12] on 33 Indian populations normalized over social groups, language families and their geographical distribution (details in **Materials and Methods** and **Supplementary Note**) and plotted the top two left singular vectors of the resulting genetic covariance matrix (**Figure 1** and **Supplementary Figures 1 and 2** for the entire dataset), with the first three PCs explaining 32%, 15% and 10% of the total variance, respectively. Along PC1, we observed a separation of TB speakers from the rest of the Indian populations. On the other hand, the IE and DR speaking populations formed a cline separated from AA speakers in PC2. Next, we computed the Pearson correlation coefficient (*r*^2^) between the top two left singular vectors (we will denote them by PC1 and PC2) of the covariance matrix and the geographic coordinates (longitude and latitude) of the samples under study. We observed *r*^2^ = 0.604 (*p* < 10^−9^) for PC1 vs. longitude and *r*^2^ = 0.065 (*p* < 10^−9^) for PC2 vs. latitude. Thus, PC1 correlates well with longitude due to the East-West cline of language families with IE and TB speakers in Northwestern and Northeastern frontiers, respectively and AA speakers dwelling in the forests of Central India between them. However, PC2 only minimally correlates with latitude, just barely picking up a previously reported North-South cline of IE and DR speakers [53] (**Figure 1B**). We note that IE and DR speakers also share significant ancestry among SGA and SGB groups (**Supplementary Figure 3**). Interestingly, in the second and third PCs, we observed further sociolinguistic stratification with SGCs distinguished from SGA and SGB within their language group (**Supplementary Figure 4**).

This weak correlation between geography and genetics in Indian context is confirmed by Mantel tests between genetic (*F*_*ST*_) and geographic distances which returned a low *r*^2^ = 0.17 (*p* = 0.0001, *Z* = 5.71) when run on the normalized data set with 33 groups. These findings are in sharp contrast with findings within the European continent [42], [26] and highlight the need for social and linguistic factors to be accounted for, as noted in prior work [7], [55], [14], [35], [9]. We performed Linear Discriminant Analysis (LDA) (**Supplementary Figures 5**) in order to gain further understanding of the relationship between genetics, geography, language and social groups in shaping the structure of the data. We run LDA on the normalized subset with the classes set as language groups (**Supplementary Figures 5A**) and then as geographic regions (**Supplementary Figures 5B**). In the LDA performed by language group, three separate clusters capturing IE social groups (SGA, SGB and SGC) appear in one axis of variation. The second axis captures the rest of the language groups again stratified by social group. In the LDA performed by geography, we see an east-west cline with TB speakers in the left and IE speakers in the right along the first discriminant. However, the second discriminant does not pick up the north-south cline as was expected, further indicating confounding by sociolinguistic groups.

### Correlation Optimization of Genetics and Geodemographics

Having shown that geography alone cannot explain the genetic structure within India, we applied COGG to explore whether integrating information on spoken language and social structure as shaped by endogamy can lead to an improved model. Indeed, solving the optimization problem that underlies COGG (see **Materials and Methods** and **Supplementary Note** for the exact formulation) and plugging in the solution, we observe a significant increase in a Pearson correlation coefficient (*r*^2^), from 0.6 to 0.93 (*p* < 10^−22^) for PC1 vs. **G** and from 0.06 to 0.85 (*p* < 10^−15^) for PC2 vs. **G**. Our results clearly show that endogamy and language families are pivotal in studying the genetic stratification of Indian populations. This is in sharp contrast to what has been seen in other parts of the world where geography is a major contributor in shaping genetic structure of populations [15], [42], [5]. Our results are statistically significant (**Supplementary Figure 6**) over 1,000 iterations with permutation of the variables related to social factors and languages (see **Supplementary Note**).

We further explored an extension of COGG in order to jointly analyze multiple PCs simultaneously and not just each component individually. To do this, we employed Canonical Correlation Analysis (CCA), a well-studied statistical technique, which maximizes the correlation between the genetic and the Geodemographic matrices by jointly finding linear combinations of the variables in each matrix. We used the top eight PCs of the genetic matrix as the results did not improve significantly, beyond that (**Supplementary Figure 7**). We note that these eight principal components capture, collectively, 89% of the variance of the genetic matrix.

Running COGG-CCA on these inputs returns a statistically significant (**Supplementary Figure 7**) *r*^2^ equal to 0.94 (which is well above the *r*^2^ = 0.74 obtained when COGG-CCA is run without including the sociolinguistic factors and significant with *p* < 10^−16^. See **Supplementary Note** for details.)

### Identifying the features that drive population structure within India

In order to formally investigate which of these nine features in the Geodemographic matrix **G** contribute more in the optimization problem posed by COGG, we used the sparse approximation framework and the Orthogonal Matching Pursuit (OMP) algorithm from applied mathematics ([41]) (see **Supplementary Note**). Running OMP on our dataset we obtain two sets of three features each, *S*_1_ and *S*_2_, for PC1 and PC2 respectively:

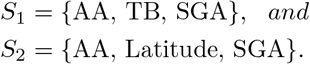

Plugging in *S*_1_ as the reduced feature space in COGG resulted in *r*^2^ = 0.92 (*p* < 10^−15^) for PC1 vs. *S*_1_ and 0.85 (*p* < 10^−12^) for PC2 vs. *S*_2_. These values capture over 99% of the correlation returned by COGG when all the features in *G* are included. Membership to the AA and TB language groups which are identified among the top significant features correspond mostly to tribal nomadic hunter gatherers dwelling in the hills and forests of Central East and North East India, respectively. Thus, the AA and TB language groups automatically capture SGC. On the other hand, membership to SGA, which is the other top significant feature that we identified, spans most of the IE and DR speakers found across Northern and Southern India. Thus, these three features appear to encompass most of the geographic, social and linguistic diversity found in the Indian subcontinent and highlight their interplay.

### Ethnic groups capturing genetic diversity across India

We developed a simple approach based on the Ridge Leverage Score (RLS) statistic ([2]) (see Equation **??** and **Supplementary Note** for details) to identify influential (from a genetic perspective) Indian populations which represent and capture the greatest portion of observed genetic diversity across India. Here, we analyzed the full pan-Indian data set of 90 populations (details in **Materials and Methods**).

The RLS statistic highlights ethnic groups in the Indian subcontinent who either are quite distinct (e.g. underwent a founder event, or practiced endogamy and maintained isolation from other groups) or populations that show signs of admixture from distinctly different language families (**Table 1**). Such populations create a mesh of complex layers of admixture across language and social barriers. We observe mostly SGB and SGC populations across all the language families in India encapsulate much of its genetic structure. Some of the highlighted populations are: (1) Great Andamanese and Jarawas from Andaman and Nicobar archipelago, which represent distinct ethnic groups and outliers with respect to mainland Indian populations (**Supplementary Figure 2B**). Great Andamanese are also linguistically divergent from Jarawa ([1]); (2) Vysyas, who underwent a founder event going back 100 generations, due to the strong imposition of endogamy ([53]); (3) Language isolates Vedda from Sri Lanka ([18]); (4) Minicoy from Lakhswadeep archipelago with strong founder effects and diverse mixture due to the archipelago being a popular destination for maritime sailors ([56]); (5) AA speaking Mundas who have Ancestral North and South Indian ancestry and an Ancestral Southeast Asian component ([61]); (6) Manipuri Brahmins (TB_SGA) who show high shared ancestry with IE_SGA as well as TB_SGC (**Supplementary Table S2**), since they are at the junction of both language families and (7) TB speaking Changpas, who are semi-nomadic pastoralits dwelling in the high altitudes of Tibet and Ladakh in India.

**Table 1.** Top ten significant ethnic groups in India capturing the genetic structure of the subcontinent as reflected by the RLS statistic (* Vysyas are classified as in between SGA and SGB [39].

| Population group | State/Territory | Language family | Social group |
| --- | --- | --- | --- |
| Great Andamenese | Andaman and Nicobar islands | Great Andamanese | SGC |
| Minicoy | Lakshwadeep islands | IE | SGB |
| Vedda | Sri Lanka | IE | SGC |
| Vysya | Andhra Pradesh | DR | SGA * |
| Palliyar | Tamil Nadu | DR | SGC |
| Munda | Madhya Pradesh | AA | SGC |
| Changpas | Jammu and Kashmir | TB | SGC |
| Manipuri Brahmins | Manipur | TB | SGA |
| Meghawal | Rajasthan | IE | SGB |
| Jarawa | Andaman and Nicobar islands | Ongan | SGC |

### Relationship between sociolinguistic groups

Our analyses using COGG clearly support the fact that language families and endogamy within social groups have played a significant role in shaping the genetic structure of the Indian subcontinent. Here, we further dissect the relationship between the endogamous social groups including the Andaman isolates [62], [38] in order to highlight the cryptic relatedness among ethnic groups that COGG posits.

The SGA populations across language groups share approximately 85% average ancestry with SGC belonging to the same language group (**Supplementary Figure 3B**). The IE and DR language groups show more homogeneity in shared ancestry than the TB and AA groups. This supports the notion that there was mixture between IE and DR speakers across SGA and SGB (**Supplementary Figure 8**) around 1,900 to 4,200 years ago ([39]) and that the caste system originated from a “classless” society, which became hierarchical with the knowledge of agriculture [31], [34].

To better illustrate the intricacies in the relationships between the social groups in India, we constructed a network of all the 90 populations across India (**Figure 2**). The network was built as we have previously described [44] based on weights that reflect shared ancestry (**Supplementary Table S2**) as computed by meta-analysis of ADMIXTURE results [3] (see **Materials and Methods** and **Supplementary Note** for details). The shared ancestry network, revealed four clusters, mostly owing to the sociolinguistic stratification.

#### IE and DR populations across social groups

A cluster of IE and DR speakers across social groups resembles a nearly complete graph with over 60% of all possible edges presented (**Figure 2**). We observe a similar pattern of strong shared ancestry in outgroup *f*_3_ statistics [47], [48] using Yorubans in Nigeria (YRI) from the 1000 Genomes phase3 dataset as the outgroup [5]. We find that IE and DR populations share more alleles with each other (**Supplementary Figure 9**) with some exceptions such as Paniyas, Kadar, and Irula (also observed in **Figure 2**).

*f*_3_ tests for signs of admixture (**Supplementary Table S3**), and meta-analyses of ADMIXTURE (**Supplementary Figure 8**) revealed similar patterns, with IE speakers showing more homogeneity across social groups than DR speakers (see also **Supplementary Note** for more details).

#### Isolation of DR speakers of Western Ghats

Few DR_SGC groups such as Kadar, Irula, Palliyar, and Paniya (which contain the lowest levels of Ancestral North Indian ancestry among Indian populations ([39])) formed a connected component, isolated from the main IE-DR cluster. They are hunter gatherer populations dwelling in the forests of Western Ghats in Southern India, isolated from the rest of the DR SGCs and very low shared ancestry with IE_SGC (**Supplementary Figure 8**).

#### AA speakers forming a clique

Almost all AA populations from Central and East India tightly cluster together with fellow Central Indian groups such as Bhunjia (IE_SGC), Gonds (DR_SGB) and Sahariya (IE_SGB).

The Gonds and Sahariyas are a candidate mosaic Indian population, which is also reflected by their location as bridge nodes in the graph. They contain high AA, DR and IE ancestry (**Supplementary Figures 8 and 9** and **Supplementary Table S2**), which can be attributed to their central location in India [21] and their long history of exogamy.

#### Clique of TB speakers connected to Great Andamanese

TB speakers from North East India forms a cluster connected to the Khasis (AA speakers residing in North East India). The cluster also contain Manipuri Brahmins (TB_SGA), who are known to have significant admixture from IE_SGA (see **Supplementary Table S3**) and Tharus [20] from Tarai region in Nepal and eastern India.

The cluster reveals interaction of TB speakers of North East India with the Great Andamanese with ~ 50% shared ancestry (**Supplementary Table S2**) as well as showing strong shared genetic drift with respect to outgroup *f*_3_ statistics (**Supplementary Figure 9**). The Great Andamanese is known to be genetically divergent from other Andamanese groups Jarawa and Onge [62], [1]. To the best of our knowledge, this is the first observed interaction of the group with rest of mainland Indian populations.

#### Isolated Andamanese groups

The other Andamanese groups Jarawa and Onge diverge from the rest of the Indian populations. This has also been shown in [62], [53], [9], [38]. They belong to the Ongan language family which has a debatable connection with Austronesian languages [11], showing divergence from all language families in mainland India.

### The mosaic of Indian sociolinguistics in the context of Eurasia

Indian populations from diverse sociolinguistic groups have different genetic affinites towards Eurasian populations [53], [39], [29], [9], [46], [40]. To understand their relationship to each group, we analyzed the outgroup *f*_3_ statistics, using YRI as the outgroup (**Supplementary Table S1C**) as done in previous studies on South Asia [6, 46]. We test the extent of allele sharing between the defined social groups of AA, DR, IE and TB speakers in India and Eurasian populations. We conduct *f*_3_ tests such as *f*_3_(*Y RI*; *X, Y*), where *X* is an Indian group and *Y* is a Eurasian group. Outgroup *f*_3_ statistics reveal European populations showing greater shared genetic drift, with the IE social groups, along with DR_SGA.

The East Asian populations have more shared drift with the TB speakers along with some affinity with AA speakers, which is in agreement with a previous study [61]. Our results clearly show a gradient of genetic drift from Siberia, then Mongolia, splitting, on one hand, towards China and North East India and, on the other hand, towards the Uygurs, Central Asia, Middle Easterners, and Europeans. This is concordant with our findings from network analysis with respect to connections with possible gateways to and from the Indian subcontinent (**Supplementary Figure 10**).

## Conclusion

India represents a country of great social and linguistic complexity. We established a quantitative deterministic and non-parametric framework called COGG, aiming to evaluate the relative contribution of language, social structure and geography in shaping the Indian gene pool. COGG resulted in a dramatic increase in the Pearson correlation coefficient (*r*^2^) between top PCs depicting genomic structure and the geodemographic factors that we investigated. We applied a feature selection algorithm to identify the most important factors shaping genomic structure in India, as well as a RLS statistic to highlight ethnic groups in India that best capture its diverse gene pool. Intriguingly, our study shows that spoken language seems to have been the major force bringing people together in India, across geographic and social barriers. This is in sharp contrast with what has been shown for other parts of the world where geography seems to have a major impact [15], [45], [5] and highlights the need for population-specific studies.

We find evidence for wide mixture across all the social groups (tribal and non-tribal) for IE speakers and across SGA and SGB for DR speakers in India. This is in concordance with a previous study [34] that indicates the origin of the caste system from a semi-nomadic society which became hierarchical with the knowledge of agriculture [31], [34]. As shown previously, this wide admixture came to an abrupt end around 1,900 to 4,200 years ago [39].

IE languages (primarily spoken in Northern India) are part of a larger language family that includes a great majority of European languages. In contrast, DR languages (primarily spoken in Southern India) are not closely related to languages outside of South Asia. Nevertheless, the earliest Hindu text (the Rig Veda, written in archaic Sanskrit) contains DR loanwords that are not found in IE languages outside the Indian subcontinent [36], [39], [67]. Further supporting the long contact between IE and DR speakers in India, our network analysis and *f*_3_ tests identify a large cluster consisting of IE and DR populations which resembles an almost complete graph with almost all pairs of populations connected to each other.

We observe a stronger shared ancestry between the Great Andamenese with TB speakers of North East India than other mainland speakers. This is the first time this observation is made based on autosomal markers and should be interpreted with caution due to small sample sizes of all the groups involved. However, a study focused on the mitochondrial haplogroup M31 showed that with the exception of M31a1 (specific to Andaman archipelago), lineages M31a2, M31b and M31c are prevalent in North East India and surrounding regions [66]. The authors concluded with time estimation that the Andaman archipelago was likely settled by modern humans from North East India *via* the land-bridge connecting Andaman archipelago and Myanmar around Last Glacial Maximum (LGM) [65], [23].

The framework we have developed and applied in order to understand genetic structure within the Indian subcontinent can be applied more broadly in order to model the interaction between different factors that may have shaped genetic diversity and population structure. The possibility to correlate genomic background to geographic, social and cultural differences opens new avenues for understanding how human history and mating patterns translate into the genomic structure of extant human populations

## Data availability

Data used in this manuscript is available from the respective corresponding authors. Code for COGG and COGG-CCA is available here: https://github.com/aritra90/COGG.

## Supplementary Material

Supplementary note, tables S1 - S4 and figures S1 - S10 are available as attachments.

## Acknowledgments

This study was supported by NSF IIS-1319280, NSF IIS-1661760, and IBM. We thank D. Reich and P. Moorjani for sharing genotypic data of 248 samples from Reich et al. (2009) [53] and 378 samples from Moorjani et al. (2013) [39]. We also thank P. P. Majumder who allowed us to use the genotypic data from 367 samples from Basu et al. (2015) [9].

## Notes

### Competing Interest Statement

The authors have declared no competing interest.

### Summary of Updates

We have introduced two new methods to dissect the genetic diversity within India. We also included more data to encompass the Andaman islanders in the analysis. Extensive f3 Outgroup analysis was done to understand the shared genetic drift experienced by sociolinguistic groups in India with respect to Eurasia.

