## Supplementary material for "Integrating linguistics, social structure, and geography to model genetic diversity within India": Supp_all.pdf

### Supplementary Note

#### Normalized data sets

We assembled 48,225 genotypes for 891 samples from 90 well-defined ethnic groups (see **Supplementary Table 1**) which were collected from various sources [1–5]. After quality control for minor allele frequency (MAF) > 0.05, missingness rate less than 0.05 and Hardy-Weinberg test statistic less than 0.001 we removed 942 markers to end up with 47,283 autosomal SNPs. We did not use the Indian samples from the 1000 Genomes [6] project because of unavailability of their geographical coordinates as well as caste and language information. Additionally, three (GIH, STU, ITU) out of the five Indian population groups in the 1000 Genomes project were collected from Indian Diaspora living in the USA (Houston) and the UK and might be biased and/or lead to gross underestimation of genetic diversity.

As the consolidated data set was put together from so many varied sources, there was an imbalance of social group and language family representation (Table A) in the samples.

| Language Groups | | | | | Social Groups | | |
|---|---|---|---|---|---|---|---|
| AA | AND | DR | IE | TB | SGA | SGB | SGC |
| 131 | 52 | 336 | 279 | 93 | 207 | 211 | 473 |

Table A: Number of samples per social and language groups in the entire consolidated data set as shown in Table S1A.

We had 16 SGA, 26 SGB, 48 SGC population groups as well as 15 AA, 3 AND, 32 DR, 29 IE and 11 TB groups respectively for the entire data set. To create the normalized data set, we removed the population group Garo from the TB data set as the social group they belong to were unknown. Thus, the resulting data set had 90 individuals from TB and we sub-sampled a

| Language Groups | | | | Social Groups | | |
|---|---|---|---|---|---|---|
| AA | DR | IE | TB | SGA | SGB | SGC |
| 92 | 93 | 94 | 89 | 107 | 43 | 218 |

Table B: Number of samples per social and language groups in the normalized data set. The normalization was done by language and geographical regions.

similar number of individuals from the other three language families. The sub-sampling was done with respect to the social group affiliation and geographical locations. As AA and TB speakers are more homogeneously located in the forests and hills of Central, East, and Northeast India, and, on the other hand, IE and DR speakers are more spread across the northern and southern India, we sampled individuals in order to guarantee a balanced representation of geographical variance. We also made sure that all social groups are equally represented in the normalized data set. This resulted in having 368 individuals sampled across 33 population groups from all over India (Table B). We created multiple normalized subsets of the original consolidated data set using the same technique. For example, as shown in Table S1B, we included Kashmiri Pandits and Kshatriya for IE_SGA in the normalized subset used in this study. However, to check robustness, we included Brahmins and Srivastava for another subset and the same was done for each sociolinguistic category. This was done to check the robustness of our results. Indeed, all our analyses returned similar results with very minor changes in the squared correlation values.

#### Correlation Optimization of Genetics and Geodemographics (COGG)

We now describe in more detail the proposed Correlation Optimization of Genetics and Geodemographics (COGG) method, which maximizes the correlation between one of the top two principal components and the geodemographic matrix, containing geographical coordinates, caste, tribe and language information. We restrict our encoding into three castes: SGA, SGB and SGCs, naming them as such instead of the conventional derogatory terms that are widely used. We noticed that the Middle castes are very similar to the Forward castes, such as Kshatriya or Brahmins, hence, we labelled both Forward and Middle castes as SGA. Although the term Backward Class (as well as Scheduled castes and Scheduled Tribes) is used by the Government of India to classify social groups which are socially and educationally disadvantaged, we chose to call them SGB.

Let $\mathbf{u}$ be the $m$-dimensional vector containing (say) the top left singular vector of the genetic covariance matrix (PC1), as computed by a software such as EIGENSTRAT [7], and let $\mathbf{G}$ denote the Geodemographic matrix, as follows:

$$\mathbf{G} = \begin{array}{c} \\ \end{array} \left[ \begin{array}{c|c|c|c|c|c|c|c|c} G_1 & G_2 & G_3 & G_4 & G_5 & G_6 & G_7 & G_8 & G_9 \\ \vdots & \vdots & \vdots & \vdots & \vdots & \vdots & \vdots & \vdots & \vdots \\ \vdots & \vdots & \vdots & \vdots & \vdots & \vdots & \vdots & \vdots & \vdots \\ Latitude & Longitude & SGA & SGB & SGC & AA & DR & IE & TB \\ \vdots & \vdots & \vdots & \vdots & \vdots & \vdots & \vdots & \vdots & \vdots \\ \vdots & \vdots & \vdots & \vdots & \vdots & \vdots & \vdots & \vdots & \vdots \end{array} \right]$$

The social groups (SGA, SGB and SGC) and Language (AA, DR, IE, TB) encoding was done as follows:

$$\textbf{Castes (or Languages)} = \begin{cases} 1, & \text{if the sample belongs to that social group (or Language)} \\ 0, & \text{otherwise} \end{cases}$$

Let $\mathbf{a}$ be the $k$-dimensional vector whose elements are $a_1 \dots a_k$ (in our case, $k = 9$). COGG solves the following optimization problem:

$$\max_{\mathbf{a}} \mathbf{Corr} \left( \mathbf{u}, \sum_{i=1}^{k} a_i \mathbf{G}_i \right). \quad (1)$$

Recalling the definition of the Pearson correlation coefficient, we can rewrite the above optimization problem as

$$\max_{\mathbf{a}} \mathbf{Corr}\left(\mathbf{u}, \sum_{i=1}^{k} a_i \mathbf{G}_i\right) = \max_{\mathbf{a}} \frac{\mathbf{u}^T(\sum_{i=1}^{k} a_i \mathbf{G}_i)}{\sqrt{\mathbf{Var}\left[\mathbf{u}\right] \mathbf{Var}\left[\sum_{i=1}^{k} a_i \mathbf{G}_i\right]}} = \max_{\mathbf{a}} \frac{\sum_{i=1}^{k} a_i(\mathbf{u}^T \mathbf{G}_i)}{\sqrt{\mathbf{Var}\left[\mathbf{u}\right] \sum_{i,j=1}^{k} a_i(\mathbf{G}_i^T \mathbf{G}_j) a_j}}. \quad (2)$$

Let $d_i = \mathbf{u}^T \mathbf{G}_i / \sqrt{\mathbf{Var}\left[\mathbf{u}\right]}$ for $i = 1 \ldots k$ and let $\mathbf{d}$ be the vector of the $d_i$'s. Also, let $\mathbf{M}_{ij} = \mathbf{G}_i^T \mathbf{G}_j$ for all $i, j = 1 \ldots k$ and let $\mathbf{M}$ be the matrix of the $\mathbf{M}_{ij}$'s. By definition, $\mathbf{M}$ is a square, symmetric positive definite matrix and hence its square root $\mathbf{M}^{1/2}$ is well-defined. We can now rewrite the above equation as

$$\max_{\mathbf{a}} \left(\mathbf{u}, \sum_{i=1}^{k} a_i \mathbf{G}_i\right) = \max_{\mathbf{a}} \frac{\mathbf{d}^T \mathbf{a}}{\sqrt{\mathbf{a}^T \mathbf{M} \mathbf{a}}} = \max_{\mathbf{a}} \frac{\mathbf{d}^T \mathbf{a}}{\|\mathbf{M}^{1/2} \mathbf{a}\|_2}.$$

To understand the last equality let $\|\mathbf{x}\|_2$ denote the Euclidean norm of the vector $\mathbf{x}$ and recall that: *(i)* since $\mathbf{M}$ is symmetric positive definite matrix, $\mathbf{M} = (\mathbf{M}^{1/2})^T \mathbf{M}^{1/2}$ and *(ii)* $\sqrt{\mathbf{x}^T \mathbf{x}} = \|\mathbf{x}\|_2$ for any vector $\mathbf{x}$, including $\mathbf{x} = \mathbf{M}^{1/2} \mathbf{a}$. Now assume that $\mathbf{M}$ is invertible and make the change of variable $\mathbf{p} = \mathbf{M}^{1/2} \mathbf{a} / \|\mathbf{M}^{1/2} \mathbf{a}\|_2$. Notice that $\mathbf{p}$ is a unit norm vector (its Euclidean norm is equal to one) and that

$$\mathbf{a} = \|\mathbf{M}^{1/2} \mathbf{a}\|_2 \mathbf{M}^{-1/2} \mathbf{p}. \quad (3)$$

Thus, we get:

$$\max_{\mathbf{p}, \|\mathbf{p}\|_2 = 1} \left(\mathbf{u}, \sum_{i=1}^{k} a_i \mathbf{G}_i\right) = \max_{\mathbf{p}, \|\mathbf{p}\|_2 = 1} \mathbf{d}^T \mathbf{M}^{-1/2} \mathbf{p}. \quad (4)$$

Using sub-multiplicativity and the fact that $\mathbf{p}$ is a unit norm vector,

$$\mathbf{d}^T \mathbf{M}^{-1/2} \mathbf{p} \leq \|\mathbf{d}^T \mathbf{M}^{-1/2}\|_2 \|\mathbf{p}\|_2 = \|\mathbf{d}^T \mathbf{M}^{-1/2}\|_2 = \sqrt{\mathbf{d}^T \mathbf{M}^{-1} \mathbf{d}}. \quad (5)$$

The last equality follows from the fact that $\|\mathbf{x}\|_2 = \sqrt{\mathbf{x}^T \mathbf{x}}$ for any vector $\mathbf{x}$. The above upper bound is true for any unit norm vector $\mathbf{p}$ and can actually be achieved by the vector $\mathbf{p}_{\max}$:

$$\mathbf{p}_{\max} = \frac{\mathbf{M}^{-1/2} \mathbf{d}}{\|\mathbf{M}^{-1/2} \mathbf{d}\|_2}.$$

Indeed, it is easy to verify that $\mathbf{p}_{\max}$ is a unit norm vector that satisfies

$$\mathbf{d}^T \mathbf{M}^{-1/2} \mathbf{p}_{\max} = \mathbf{d}^T \mathbf{M}^{-\frac{1}{2}} \frac{\mathbf{M}^{-1/2} \mathbf{d}}{\|\mathbf{M}^{-1/2} \mathbf{d}\|_2} = \frac{\mathbf{d}^T \mathbf{M}^{-1} \mathbf{d}}{\sqrt{\mathbf{d}^T \mathbf{M}^{-1} \mathbf{d}}} = \sqrt{\mathbf{d}^T \mathbf{M}^{-1} \mathbf{d}}.$$

Thus, from eqn. (5), it follows that $\mathbf{p}_{\max}$ is a maximizer for the optimization problem of eqn. (4). If we let

$$\mathbf{a}_{\max} = \mathbf{M}^{-1} \mathbf{d},$$

it is easy to see that the above values for $\mathbf{a}_{\max}$ and $\mathbf{p}_{\max}$ satisfy

$$\mathbf{a}_{\max} = \|\mathbf{M}^{1/2} \mathbf{a}_{\max}\|_2 \mathbf{M}^{-1/2} \mathbf{p}_{\max},$$

as stipulated by the change of variables from eqn. (3), and thus $\mathbf{a}_{\max}$ is a maximizer for COGG.

Plugging in the solution for $\mathbf{a}$, COGG revealed a squared Pearson Correlation coefficient $r^2 = 0.93$ for PC1 vs $\mathbf{G}$ and $r^2 = 0.85$ for PC2 vs $\mathbf{G}$. These values represent a manyfold increase from the original correlation values of $r^2 = 0.6$ for PC1 vs $\mathbf{G}'$ and $r^2 = 0.06$ for PC2 vs $\mathbf{G}'$, where, $\mathbf{G}'$ is the matrix $G$ without the sociolinguistic features, containing only the geographical coordinates. This highlights that geography is not enough as a feature to understand the genetic structure of the Indian populations.

In order to use the pre-defined caste and language group affiliations and understand the most significant variables among social structure, language, and geography we tried a different encoding scheme where we encoded social groups and languages as two variables instead of the previously used three variables (SGA,

SGB, SGC) for castes and four variables (AA, DR, IE, TB) for languages. Now, we assigned 1, 2, and 3 for SGA, SGB and SGC groups, respectively, in the social category and similarly 1, 2, 3, and 4 for AA, IE, DR, TB affiliations in the language variable. This removes the sparsity induced by the zero-one indicator variable encoding as previously used. Running COGG with this encoding resulted in values of $r^2$ equal to 0.79 for PC1 and 0.82 for PC2, respectively. We observe that the value of $r^2$ for PC1 shows a decrease from 0.92 (when the previous encoding was used) to 0.79, whereas for PC2 it shows a smaller decrease. To understand the loss of $r^2$ for PC1, we ran the feature selection algorithm (discussed in the next section) and noticed that the most significant variable is longitude followed by the consolidated caste variable. Previously, we noticed that the most significant variables were AA, TB and the SGA. However, now, AA and TB are both encoded in a single language variable and we see that we lose some interpretability of the individual variables which capture the genetic variation of the Indian subcontinent.

We also investigated whether the values returned by COGG are statistically significant. We performed 1,000 iterations with randomly permuted values of the columns related to caste and language encoding in $\mathbf{G}$. We do not permute the columns corresponding to the geographical coordinates in order to maintain a baseline for the comparison. We randomly permuted the rows (individuals) corresponding to the seven columns (variables related to castes and language affiliations) in $\mathbf{G}$ and in each iteration we run COGG to find the optimal $\mathbf{a}_{\max}$ and the respective $r^2$ value. We find that the random permutations return a maximal value of $r^2$ equal to 0.6422 for PC1 and 0.1679 for PC2 (Figure S6). This is a minor increase from 0.6 and 0.06 respectively for PC1 and PC2, clearly indicating the importance of the caste and language encoding in $\mathbf{G}$.

Prior work attempted to disentangle the effects of non-genetic variables such as geography, linguistics, subsistence, social or ecological factors from the genetic variables captured by the top principal components. One such study [8] regressed the top 20 PCs computed from the genotypes of the Khoe-San populations with various combinations of geographic, linguistic and subsistence covariates, and used cross-validation scores to understand which non-genetic variable can predict the observed genetic patterns. They observed that languages improve the predictive capacity of a model that includes only geography in the sub-Saharan and the Southern African data set. This is similar to the intuition of COGG, which provides a conceptually straightforward model to do an in-depth study to account for the factors within the broad generic non-genetic factors, such as which language and social group explain most of the genetic variation captured by the top principal components. Also, in addition, we do a feature selection procedure to obtain the most significant variables in the geodemographic matrix, unlike previous studies. Another study [9] employed a Bayesian framework to isolate ecological factors from geographic distances. Broadly, COGG tries to achieve the same goal, but it provides the ease of use in this setting, where one can just encode the environmental and ecological factors as covariates and solve the underlying optimization problem to obtain the maximum correlation. Along with this, it is easier to comprehend, as it is closer to a linear regression setting.

#### Canonical Correlation Analysis (CCA)

Finally, there is no mathematical reason to restrict COGG to the top two principal components and corresponding singular vectors (PC1 and PC2) of the genetic similarity covariance matrix. Prior work has exclusively focused on studying the correlation between longitude and latitude and the top two principal components; COGG goes beyond this by adding geodemographic features to study more general correlations. Our next method applies Canonical Correlation Analysis (CCA, introduced in [10]) to simultaneously study the correlation between the top $q$ Principal Components (where $q$ is a user-defined parameter) and the geodemographic matrix $\mathbf{G}$. CCA extracts linear components that capture correlations between two input datasets, in a manner analogous to PCA. From a statistical point of view, CCA extracts directions of maximal "correlation" between a pair of datasets represented by matrices. From a linear algebraic point of view, CCA measures the similarities between the subspaces spanned by the columns of each of the two datasets, represented by matrices [11]. In our case, we extend the optimization problem of eqn. (1) to identify the maximal correlation to include the matrix of top $q$ principal components denoted as $\mathbf{U} \in \mathbb{R}^{m \times q}$ for $m$ individuals and $\mathbf{G}$, the geodemographic matrix as described earlier. We obtain $\mathbf{U}$ by considering the top $q$ left singular vectors of the genetic covariance matrix of the normalized subset. Formally, we define the following optimization problem, which we call COGG-CCA:

$$\max_{\mathbf{a},\mathbf{b}} \mathbf{Corr}\left(\sum_{j=1}^{q} b_j \mathbf{U}_j, \sum_{i=1}^{k} a_i \mathbf{G}_i\right), \tag{6}$$

where $\mathbf{b}$ is a $p$-dimensional vector whose entries are the $b_j$'s and $\mathbf{a}$ is a $k$-dimensional vector whose entries are the $a_i$; $\mathbf{U}_j$ and $\mathbf{G}_i$ represent the $j$-th and $i$-th column of $\mathbf{U}$ and $\mathbf{G}$ as column vectors. Solving COGG-CCA analytically dates back to the work of [10] and allows us to obtain the following closed form solution for the vectors $\mathbf{a}$ and $\mathbf{b}$, the unknown coefficient vectors associated with the matrices $\mathbf{G}$ and $\mathbf{U}$, respectively.

Let $\mathbf{\Sigma}_{UU} = \mathbf{Cov}\left[U, U\right]$, $\Sigma_{GU} = \mathbf{Cov}\left[G, U\right]$, and $\Sigma_{GG} = \mathbf{Cov}\left[G, G\right]$ denote three covariance matrices and construct

$$\mathbf{\Sigma} = \mathbf{\Sigma}_{GG}^{-1/2} \mathbf{\Sigma}_{GU} \mathbf{\Sigma}_{UU}^{-1/2}.$$

Then, $\mathbf{a}$ is the top right singular vector of the matrix $\mathbf{\Sigma}$ and $\mathbf{b}$ is the top left singular vector of $\mathbf{\Sigma}$; it is well-known that the maximum correlation coefficient is equal to the largest singular value of the matrix $\mathbf{\Sigma}$. Applying COGG-CCA on our data, we obtain the maximum value $r^2 = 0.94$ for $q = 8$. To check for statistical significance of COGG-CCA, we first formed the baseline of $r^2$ by just including the geographical coordinates in the Geodemographic matrix $\mathbf{G}$. This resulted in $r^2 = 0.74$. Next, we permuted the features in both the matrices, $\mathbf{G}$ and $\mathbf{U}$, respectively which resulted in a very small increase from the baseline with $r^2 = 0.76$, whereas COGG-CCA, even with smaller values of $q$ resulted in very high $r^2$ (Supplementary Figure S5).

#### Feature selection using Orthogonal Matching Pursuit (OMP)

We used a greedy feature selection algorithm described in [12] to select features in the Geodemographic matrix $\mathbf{G} \in \mathbb{R}^{m \times k}$ containing $m$ individuals and $k$ demographic features. The precise algorithm is described below.

**Algorithm 1** OMP Algorithm for Feature Selection

**Input:** matrix $\mathbf{G}$, column vector $\mathbf{U} \in \mathbb{R}^m$, $\epsilon > 0$
**Output:** matrix $\mathbf{C} \in \mathbb{R}^{m \times p}$ which has columns of $\mathbf{G}$ with indices in $\tau$, $|\tau| = p$, $p < k$
$\tau \leftarrow \phi$; $r \leftarrow 0$; $\mathbf{U}^{(0)} \leftarrow \mathbf{U}$; $\mathbf{G}^{(0) \leftarrow \mathbf{G}}$;$\mathbf{C} \leftarrow \phi$
**while** $\|\mathbf{U}^{(r)}\|_2 > \epsilon$ **do**
  **for** $i \in \{1, 2, \cdots, k\} - \tau$ **do**
    choose $\mathbf{i}$ corresponding to maximum $\boldsymbol{corr}\left(\mathbf{U}^{(r)}, \mathbf{G}_i^{(r)}\right)$
  **end for**
  $\tau \leftarrow \tau \cup \{i\}$; $\mathbf{V} \leftarrow \mathbf{G}_i^{(r)}$
  remove column $i$ from $\mathbf{G}^{(r)}$ to form $\mathbf{G'}^{(r)}$
  project $\mathbf{G'}^{(r)}$ onto the subspace orthogonal to $\mathbf{V}$, i.e., $\mathbf{G}^{(r+1)} \leftarrow \mathbf{G'}^{(r)} - \left(\mathbf{V}\mathbf{V}^{\dagger}\right) \mathbf{G'}^{(r)}$
  project $\mathbf{U}^{(r)}$ onto the subspace orthogonal to $\mathbf{V}$, i.e., $\mathbf{U}^{(r+1)} \leftarrow \mathbf{U}^{(r)} - \left(\mathbf{V}\mathbf{V}^{\dagger}\right) \mathbf{U}^{(r)}$
  $r \leftarrow r + 1$
**end while**
$\mathbf{C} \leftarrow \mathbf{G}_{\tau}$

#### Ridge Leverage Scores

We start with the definition of the *statistical leverage scores* of a matrix.

**Definition 1** Given an arbitrary $m \times n$ matrix $\mathbf{A}$ with $m > n$, let $\mathbf{U}$ denote the $n \times d$ matrix consisting of the $d$ *left singular vectors* of $\mathbf{A}$ and let $U_{i*}$ denote the $i^{th}$ row of the matrix $\mathbf{U}$ as a *row vector*. Then, the statistical leverage scores of the rows of $\mathbf{A}$ are given by

$$\ell_i = \|\mathbf{U}_{i*}\|_2^2$$

Classical leverage scores quantify the importance of each column $i$ for the range space of the data matrix $\mathbf{A}$. They are widely used in regression problems, outlier detection and randomized matrix algorithms. They are used to select import features from an under-determined system. To address instability issues in a regression, ridge regression is performed and an extension of this notion of *classical* leverage scores to a ridge regression setting is known as *ridge leverage scores*. It is defined as follows.

**Definition 2** The ridge leverage score ${\tau_i}^{\lambda}(\mathbf{A})$ is defined as,

$${\tau_i}^{\lambda}(\mathbf{A}) = \left(\mathbf{A}\mathbf{A}^\top \left(\mathbf{A}\mathbf{A}^\top + \lambda \mathbf{I}_n\right)^{-1}\right)_{ii}$$

where $\lambda > 0$ is the regularization parameter. Further simplifying, the row ridge leverage scores boils down to the following,

$$\begin{aligned}
{\tau_i}^{\lambda}(\mathbf{A}) &= \left(\mathbf{A}\mathbf{A}^\top \left(\mathbf{A}\mathbf{A}^\top + \lambda \mathbf{I}_n\right)^{-1}\right)_{ii} \\
&= \left(\mathbf{U}\boldsymbol{\Sigma}\mathbf{V}^\top\mathbf{V}\boldsymbol{\Sigma}^\top\mathbf{U}^\top(\mathbf{U}\boldsymbol{\Sigma}\mathbf{V}^\top\mathbf{V}\boldsymbol{\Sigma}^\top\mathbf{U}^\top + \lambda\mathbf{I}_n)^{-1}\right)_{ii} \\
&= \left(\mathbf{U}\boldsymbol{\Sigma}^2\mathbf{U}^\top(\mathbf{U}\boldsymbol{\Sigma}^2\mathbf{U}^\top + \lambda\mathbf{U}\mathbf{U}^\top)^{-1}\right)_{ii} \\
&= \left(\mathbf{U}\boldsymbol{\Sigma}^2(\boldsymbol{\Sigma}^2 + \lambda)^{-1}\mathbf{U}^\top\right)_{ii} \\
&= \left(\mathbf{U}\boldsymbol{\Sigma}_\lambda\mathbf{U}^\top\right)_{ii}
\end{aligned}$$

For the above, we have assumed that $\mathbf{A}$ has full row rank as $d >> n$. Therefore the thin SVD of $\mathbf{A}$ is $\mathbf{U}\boldsymbol{\Sigma}\mathbf{V}^\top$, where $\mathbf{U} \in \mathbb{R}^{n\times n}$, $\mathbf{V} \in \mathbb{R}^{d\times n}$ and $\boldsymbol{\Sigma} \in \mathbb{R}^{n\times n}$ whose diagonal elements are the singular values of $\mathbf{A}$. For the above simplification we have used the fact that $\mathbf{U}$ and $\mathbf{V}$ are orthogonal matrices with orthonormal columns hence, $\mathbf{U}\mathbf{U}^\top = \mathbf{U}^\top\mathbf{U} = \mathbf{I}_n$ and $\mathbf{V}\mathbf{V}^\top = \mathbf{V}^\top\mathbf{V} = \mathbf{I}_d$ and their inverse is equal to transpose. Also, from above, $\boldsymbol{\Sigma}_\lambda \in \mathbb{R}^{n\times n}$ and the $i^{th}$ diagonal entry of it is defined as,

$$(\boldsymbol{\Sigma}_\lambda)_{ii} = \sqrt{\frac{\sigma_i^2}{\sigma_i^2 + \lambda}},\ i = \{1, 2, \cdots, n\} \tag{7}$$

Thus, we can write the row ridge leverage scores as,

$$\tau^{\lambda}(\mathbf{A}) = \|\mathbf{U}\boldsymbol{\Sigma}_\lambda\|_2^2$$

Armed with this definition we devise the algorithm to calculate the RLS statistic.

**Algorithm 2** Row Ridge leverage score algorithm

**Input:** A matrix, $\mathbf{A} \in \mathbb{R}^{m\times n}$
**Output:** $\tau^{\lambda}(A) \in \mathbb{R}^{m\times 1}$
$\mathbf{B} = \mathbf{A}\mathbf{A}^\top$
Compute thin SVD of $\mathbf{B} = \mathbf{U}\boldsymbol{\Sigma}\mathbf{V}^\top$
Choose $\lambda = \text{mean}\{\sigma_1, \sigma_2, \cdots, \sigma_n\}$ where $\sigma_i$ is the $i^{th}$ diagonal element of $\boldsymbol{\Sigma}$
Compute $(\boldsymbol{\Sigma}_\lambda)$ as defined in 7
Compute $\tau^{\lambda}(\mathbf{A}) = \|\mathbf{U}\boldsymbol{\Sigma}_\lambda\|_2^2$
Return the vector $\tau^{\lambda}(\mathbf{A})$

We obtain the row ridge leverage scores in this manner for the respective mean-centered genotype matrix consisting of $m$ individuals and $n$ SNPs and the Geodemographic matrix (described earlier). Thereafter, we compute the additive ridge leverage score per population as described in the Methods.

Running COGG with the significant ethnic groups as shown in Table 1 on 90 pan-Indian populations further confirmed the importance of of these populations in shaping Indian genetics. The $r^2$ value between geographical coordinates and the PCs came out to be 0.21 for PC1 and 0.08 for PC2, when ran with populations from Table 1. COGG was run for the same populations and the values returned were $r^2 = 0.853$ for PC1 and the geodemographic matrix $\mathbf{G}$ and $r^2 = 0.794$ for PC2 and $\mathbf{G}$. Thus, COGG returns very high correlations using only the populations selected using the RLS statistics, capturing most of the variance reflected by the top PCs of the genetic matrix.

#### Estimating population admixture

We applied ADMIXTURE on the three data sets namely, the entire pan-Indian data set, the normalized Indian data set and the Eurasian data set just like we did for PCA. ADMIXTURE on the entire Indian data set (Figure S3A), with all populations, revealed that the groups SGB and SGCs for AA and TB, along with some DR_SGB and SGCs (such as Paniyas, Kadar and Irulas) show divergence from DR_SGA and IE populations. This is replicated in the ADMIXTURE output of the normalized Indian data set. When applied on the Eurasian data set, the IE and DR SGAs, along with IE_SGB and SGCs cluster together with most Northwestern Frontier populations. The TB_SGC and SGA show signs of admixture from the Chinese populations. Some Middle Eastern populations and Caucasians share the same ancestral components as the IE and DR SGA. The European populations seem to be sharing very small amount of ancestral components with the IE and DR speaking groups. To investigate further and quantify the shared ancestry between these populations we employed a quantitative meta-analysis of ADMIXTURE which was first developed in [13].

We now describe in more detail our quantitative analysis of ADMIXTURE's output. Given a target population X and reference populations Y, Z, etc., we were interested in quantifying the amount of ancestry of population X that is captured by populations Y, Z, etc. Towards that end we devised a new approach to quantitatively analyze the output of ADMIXTURE. Recall that ADMIXTURE, for a particular value of $K$, will represent each sample using $K$ coordinates. Thus, for a particular value of $K$ and for a particular population Y with $n$ samples, we can represent the output of ADMIXTURE for this population as an $n$-by-$K$ table. Then, for each reference population Y, we summarize this $n$-by-$K$ matrix using its top right singular vector only; in all our analyses, the top singular value corresponding to the top right singular vector captured at least 80% of the reference population variance as represented by ADMIXTURE. Let $v_Y$ be the top right singular vector (a $K$-dimensional vector) for population Y; similarly, let $v_Z$ be the top right singular vector (a $K$-dimensional vector) for population Z, etc. Now that we have represented the ADMIXTURE output for each population as a $K$-dimensional signature vector, we can apply standard vector space calculus in order to answer our original question: how much of the ancestry of population X is captured by population Y, or population Z, etc. More specifically, in order to compute the percentage of the ancestry of population X that is captured by population Y, we compute the percentage of the norm of $V_X$ that is captured (in projection sense) by $v_Y$. Formally, we compute

$$\frac{\|V_X - v_Y \cdot v_Y^{\dagger} \cdot V_x\|_F}{\|V_X\|_F}$$

which returns a value between zero and one. In the above, $V_X$ denotes the m-by-K matrix representing the m samples of population X with respect to the K coordinates returned by ADMIXTURE. The notation $v_Y^{\dagger}$ indicates the pseudoinverse of the vector $v_Y$, which is equal to the transpose of the vector $v_Y$, suitably normalized. It is also worth noting that the norm used in the above equation is the standard matrix Frobenius norm. In order to quantify the amount of ancestry of population X that is captured by both populations Y and Z, we form the K-by-2 matrix $V = [v_Y v_Z]$ whose columns are the vectors $v_Y$ and $v_Z$ and we compute

$$\frac{\|V_X - V \cdot V^{\dagger} \cdot V_x\|_F}{\|V_X\|_F}$$

In the above equation, $V^{\dagger}$ denotes the pseudoinverse of the matrix $V$; the matrix $VV^{\dagger}$ is a projector on the subspace spanned by the column space of $V$. Thus, we basically extract from the matrix $V_X$ the part of $V_X$ that is captured by the (subspace spanned by the) vectors $v_Y$ and $v_Z$.

The meta-analysis when applied on the pan-Indian data set (891 individuals; 90 populations) showed that AA_SGC share a small amount of ancestry with other IE and DR tribal speakers ( 19%), whereas TB_SGC are completely isolated (Figure S3B). DR_SGC show divergence from other populations, which is due to a few tribal populations such as Irula, Kadar, and Paniyas (as pointed out in Figure S2A). We investigate this further when we apply the meta-analysis on the entire Indian data set (Supplementary Table S2) and study the meta-analysis of each population group. The most significant observation is that IE and DR populations across their caste affiliations (except DR_SGC) cluster together, showing high shared ancestry among the SGA and SGB. The IE tribals also share very high ancestry with the IE and DR Forward and Backward Castes. This supports the autochthonous origin of the caste system in India.

Applying the meta-analysis of ADMIXTURE on the Eurasian data along with the normalized Indian data set shows that the IE and DR castes, along with the TB_SGA, share significant amount of ancestry with Northwestern Frontier provinces (maximum in IE_SGA, who share close to 80%), which is further validated by $f_3$ statistics. The TB_SGC and SGA share approximately 94% and 68% ancestry with the Chinese populations, as well as with Mongolia. The Uygurs, along with the whole of Central Asia seems to share a small amount of ancestry with the IE populations across social groups, as well as with DR_SGA. We see similar trends in Turkey, Caucasia and European populations, sharing more ancestry with IE and DR SGAs. These populations also share close to 20% ancestry with IE_SGCs. This shows that with the spread of IE languages as shown in Figure 5B the Forward Castes and some tribes have been in touch with the migrating populations who followed the path from Siberia and Mongolia through Central Asia and Northwestern Frontier provinces. We validate these findings with $f_3$ statistics and TreeMix analyses.

#### Linear Discriminant Analysis

The genotype score value was assigned as the sum of a value of 0 for the major allele and 1 for the minor allele for each strand. The counts for each genotype out of $N$ samples are $n_{00}$ for homozygous major allele, $n_{01}$ for a heterozygous genotype, and $n_{11}$ for the homozygous minor allele. The total score across $N$ samples will be $1 \cdot n_{01} + 2 \cdot n_{11}$, with the average being $\bar{s} = (n_{01} + 2 \cdot n_{11})/(2N)$. The average squared score is $\bar{s^2} = (n_{01} + 4 \cdot n_{11})/(2N)$, so the variance is $\mathbf{Var}[s] = \bar{s^2} - (\bar{s})^2$. Scores assigned to each genotype are scaled to be $(s - \bar{s})/\sqrt{\mathbf{Var}[s]}$. In the case of Hardy-Weinberg equilibrium, this reduces to a form proportional to that employed in Eigenstrat [7]. This adjustment was applied to PCA computations performed for comparisons with LDA in this study, as well as in the normalization of the LDA input scores.

We maintain a matrix, $d \in \mathcal{R}^{N \times D}$, where $N$ rows represent the individuals and $D$ columns represent a genotype score. There are $G$ groups of populations and each group has $p$ individuals. The matrix $d$ is indexed as $d_{gpi,k}$, where $p \in g$ $(g \in G)$ and $i \in p$, each with a vector of genotype scores indexed by $k \in D$. We define, $n_{gp} = |p|$ for $p \in g$, and $n_{g.} = \sum_{p \in g} n_{pg}$, then $N = \sum_{g \in G} n_{g.}$, the data are decomposed into components $d_{gpi,k} = x_{...,k} + x_{g..,k} + x_{gp.,k} + x_{gpi,k}$ such that $\sum_{i \in p} x_{gpi,k} = 0$, $\sum_{p \in g} n_{gp} x_{gp.,k} = 0$, and $\sum_{g \in G} n_{g.} x_{g..,k} = 0$.This produces a hierarchic decomposition of the variations among groups and populations similar to AMOVA [14], but each population and group is weighted by the number of samples they contain. Their values are determined from $x_{...,k} = N^{-1} \sum\limits_{g \in G, p \in g, i \in p} d_{gpi,k}$, $x_{g..,k} = n_{g.}^{-1} \sum\limits_{p \in g, i \in p} d_{gpi,k} - x_{...,k}$, $x_{gp.,k} = n_{gp}^{-1} \sum\limits_{i \in p} d_{gpi,k} - x_{...,k} - x_{g..,k}$, and $x_{gpi,k} = d_{gpi,k} - x_{...,k} - x_{g..,k} - x_{gp.,k}$.

The total covariance is

$$\begin{aligned} c_{k',k} &= N^{-1} \sum_{g \in G, p \in g, i \in p} (d_{gpi,k'} - x_{...,k'})(d_{gpi,k} - x_{...,k}) \\ &= N^{-1} \sum_{g \in G} n_{g.} x_{g..,k'} x_{g..,k} + N^{-1} \sum_{g \in G, p \in g} n_{gp} x_{gp.,k'} x_{gp.,k} + N^{-1} \sum_{g \in G, p \in g, i \in p} x_{gpi,k'} x_{gpi,k}. \end{aligned}$$

Then $N^{-1} \sum\limits_{g \in G, p \in g, i \in p} x_{gpi,k'} x_{gpi,k}$ represents the variation within populations, $N^{-1} \sum\limits_{g \in G, p \in g} n_{gp} x_{gp.,k'} x_{gp.,k}$ represents the variation between populations within groups, and $(S_B)_{k'k} = N^{-1} \sum\limits_{g \in G} n_{g.} x_{g..,k'} x_{g..,k}$ represents the variation between groups. The total variation within groups is:

$$(S_W)_{k'k} = N^{-1} \sum_{g \in G, p \in g} n_{gp} x_{gp.,k'} x_{gp.,k} + N^{-1} \sum_{g \in G, p \in g, i \in p} x_{gpi,k'} x_{gpi,k}$$

.

While this could be evaluated for each individual SNP by choosing $k = k'$ and probing those, it is desirable to find combinations of SNPs that are most informative of the genetic differences among groups. Those combinations may be expressed in terms of vectors $\hat{u}$ with components $\hat{u}_k$. Then the projections on the $x$'s would have the form $\sum\limits_{k \in [D]} x_{gpi,k} \hat{u}_k$. Along these projections, it is possible to write a ratio expressing the variation between groups vs within groups as $f(\hat{u}) = \frac{\hat{u}^T S_B \hat{u}}{\hat{u}^T S_W \hat{u}}$ [15,16]. Identifying $v = S_W^{1/2} \hat{u}$, this becomes $f(\hat{u}(\hat{v})) = \hat{v}^T S_W^{-1/2} S_B S_W^{-1/2} \hat{v}$. This yields stationary values where $\hat{v}$ are eigenvectors of $S_W^{-1/2} S_B S_W^{-1/2}$, with eigenvalues directly corresponding to $f$.

The genetic associations identified by the $\hat{u}$ were tested by comparing the $f$'s computed from the population groups to those obtained for samples randomly permuted among the groups. Another caveat is that the largest eigenvalues of $S_W^{-1/2}$ correspond to the smallest eigenvalues of $S_W$. Yet, these are the most sensitive to sampling variation, genotyping errors, and cumulative computational errors. Further, the smallest, most error prone eigenvalues in $S_W$ tend to dominate $S_W^{-1/2}$, as well as $f$'s, even though they do not carry useful information. We apply a threshold for a ratio between eigenvalues of $S_W$ between maximum and threshold, yielding a reciprocal square root ratio for included eigenvalues and eigenvectors in constructing $S_W^{-1/2}$. This restricts the computation to the subspace operationally spanned (or explored) by $S_W^{-1/2}$.

In general, $d$ will be $N \times D$ dimensional with $S_B$ and $S_W$ being $D \times D$. Eigenvector computational space requirements for these tends to be prohibitive. Further, $d$ will span an $N \ll D$ dimensional space. In the singular value decomposition $d = USV^T$ where $V$ is orthonormal, then $dd^T = US^2U^T$, with $S^2$ diagonal. Since $dd^T$ is symmetric, $U$ is also orthonormal. Once $U$ and $S$ were determined by diagonalization of $dd^T$, $V = d^T US^{-1}$. $V$ then represents a basis of $N$ orthogonal $D$ dimensional vectors. In that basis, $dV = US$ and $V^T S_B V$ and $V^T S_W V$ are $N \times N$ matrices. Computations of $f$ were performed in this basis.

We applied LDA on the normalized Indian data set, to look further into the geography-social group-language interplay which was revealed by COGG. We first computed LDA with the supervised regional groups such as 'North','South','East', 'Northeast','Central-East' and 'Central'. The first two discriminants (Figure S5A) reveals the stratification by the geographical locations of the individuals under study. There is a clear gradient from right to left, where it starts with groups such as Meghawals, Kashmiri Pandits, who are in the northwest, followed by the Central and Southern groups. Here, we see few clusters being dominated by the castes, although LDA was ran with geographical groups. The Bhils sit closely with AA speakers, justifying their central location in the country. The other end of the gradient is occupied by the TB speakers, with Manipuri Brahmins being closer to the IE and DR groups.

Next, LDA was ran on the same data set, but now the groupings were by language affiliations of the individuals. The groups were 'IE-SGA','IE-SGB', 'IE-SGC', etc. for all languages and castes. The top two discriminants when plotted against each other, revealed a very strong evidence on the langauge-caste interplay, as pointed out in the selected features from COGG. In Figure S3b, separate clines appear from right to left, with the first cline of IE forward castes, followed by DR and TB forward castes. Thereafter, the IE and DR backward castes appear, followed by the AA and DR SGCs and finally IE SGCs. This clearly show the genetic stratification influenced by caste groups and then language groups within the caste groups. Thus, we see a two-layer stratification, when LDA was run with the language-caste groups.

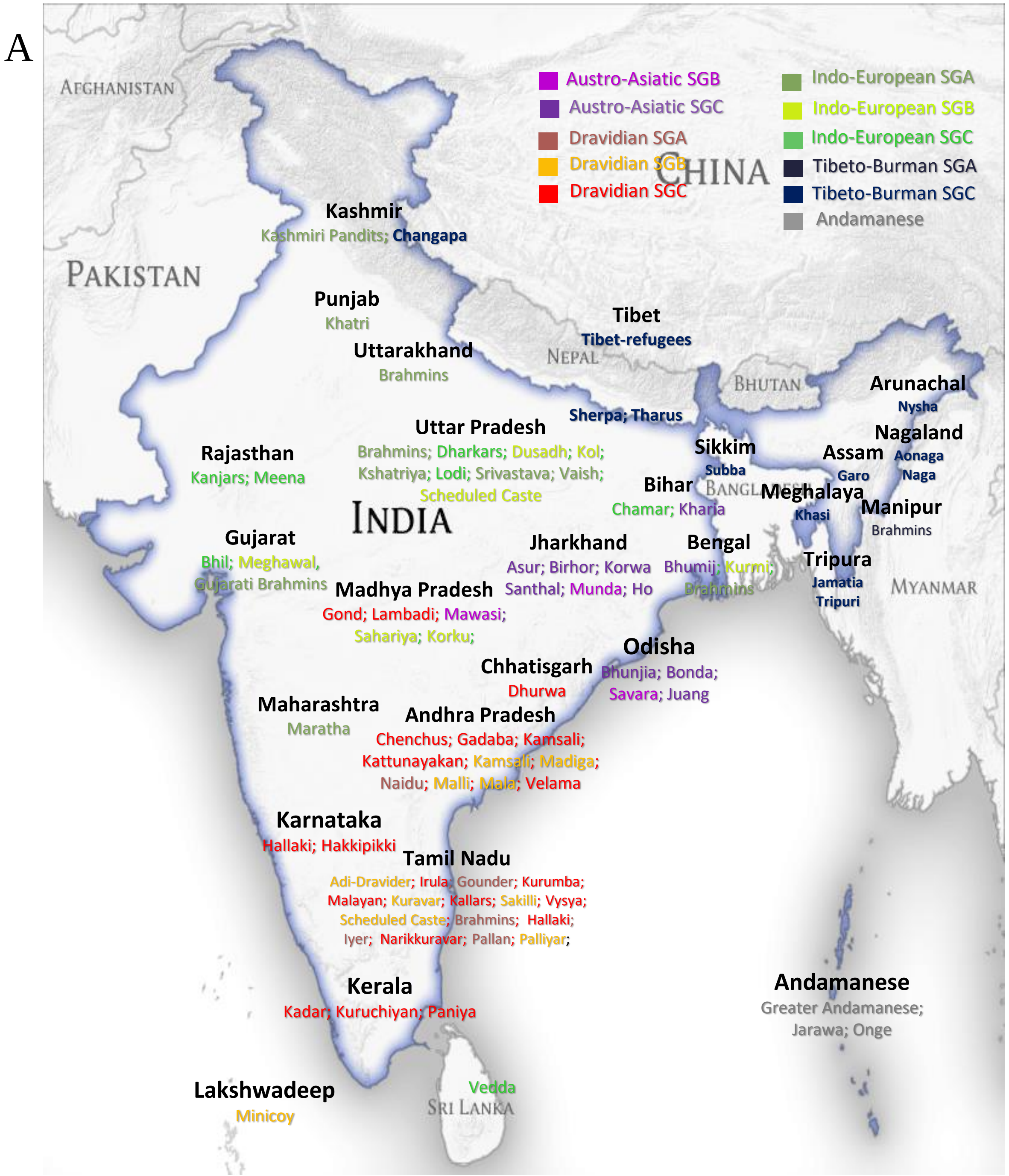


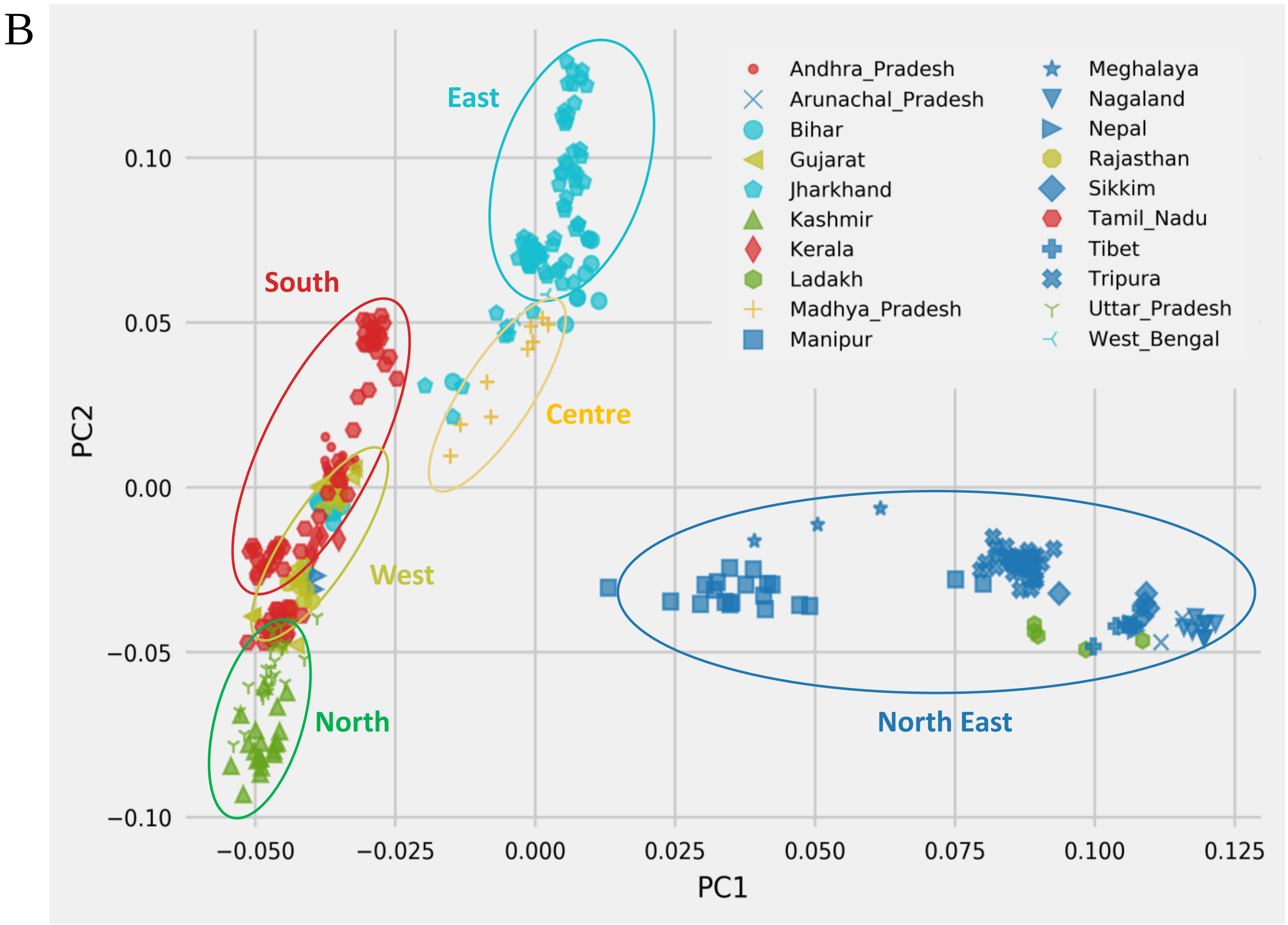


**Fig S1:** (A) Location of origin of Indian samples, grouped by geographic state and colored by sociolinguistic affiliation.

(B) Top two PCs extracted from the normalized data set consisting of 368 individuals, genotyped on 47,283 SNPs marked by geographic states and colored by geographic regions (North: Green; West: Olive; South: Red; Centre: Yellow; East: Blue and North East: Indigo) show that the top two PCs have very low correlation with geography.

**Fig S2: Population Structure of Indian populations**

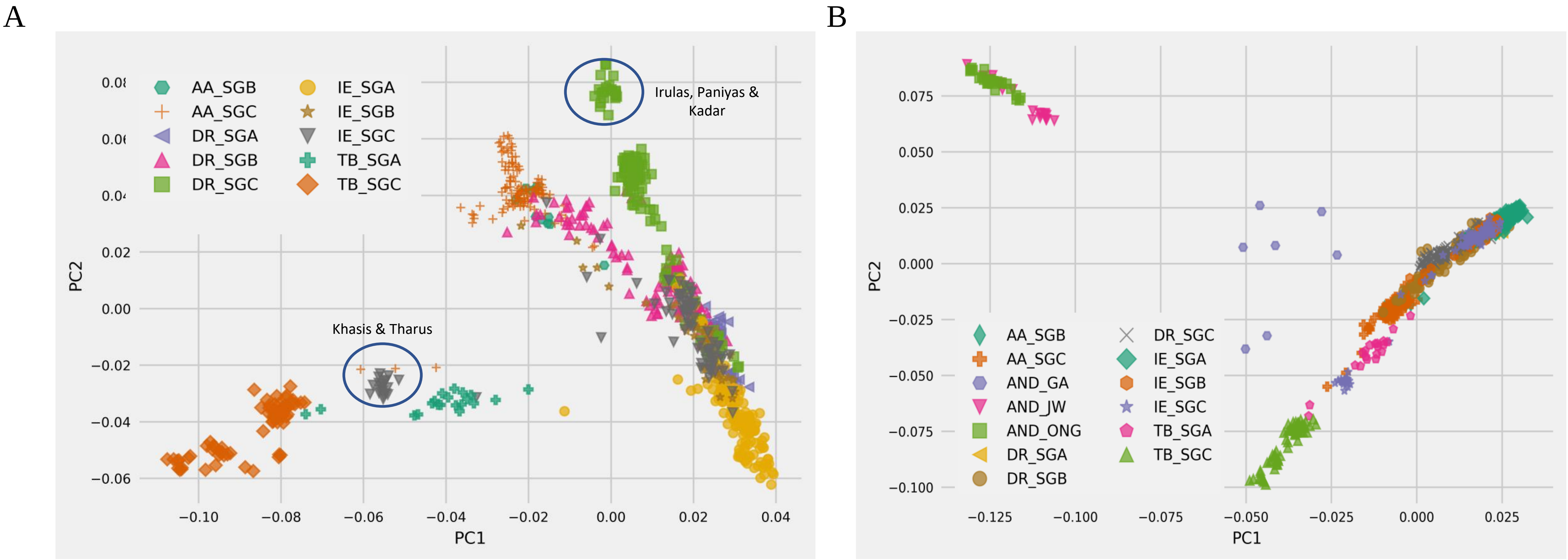


**Fig S2:** (A) PCA plot of all mainland Indian samples (839 individuals). We note that the formation of the clusters is primarily dominated by language groups, with TB_SGC and TB_SGA forming a cluster with Khasis (AA_SGC) and Tharus (IE_SGC) showing signs of admixture. The IE and DR speakers form a cline with a gradient of social groups within, IE_SGA and DR_SGC occupying the ends of the cline. We also observe that the Irulas, Paniyas, Kurumba and Kadars show divergence from other Dravidian tribal populations.

(B) ALL 891 Indian samples (including outliers from Andaman islands) projected as the top two PCs. In presence of outliers, we observe a cline for mainland Indian populations and an outlier cluster of Ongan language speaking Jarawa (AND_JW) and Onge (AND_ONG). However, the Great Andamanese (AND_GA) lies near the mainland Indian populations. Proportions of variance explained for the top 3 PCs are 33%, 22% and 12.7%,

#### Fig S3A: Admixture ancestry of Indian populations

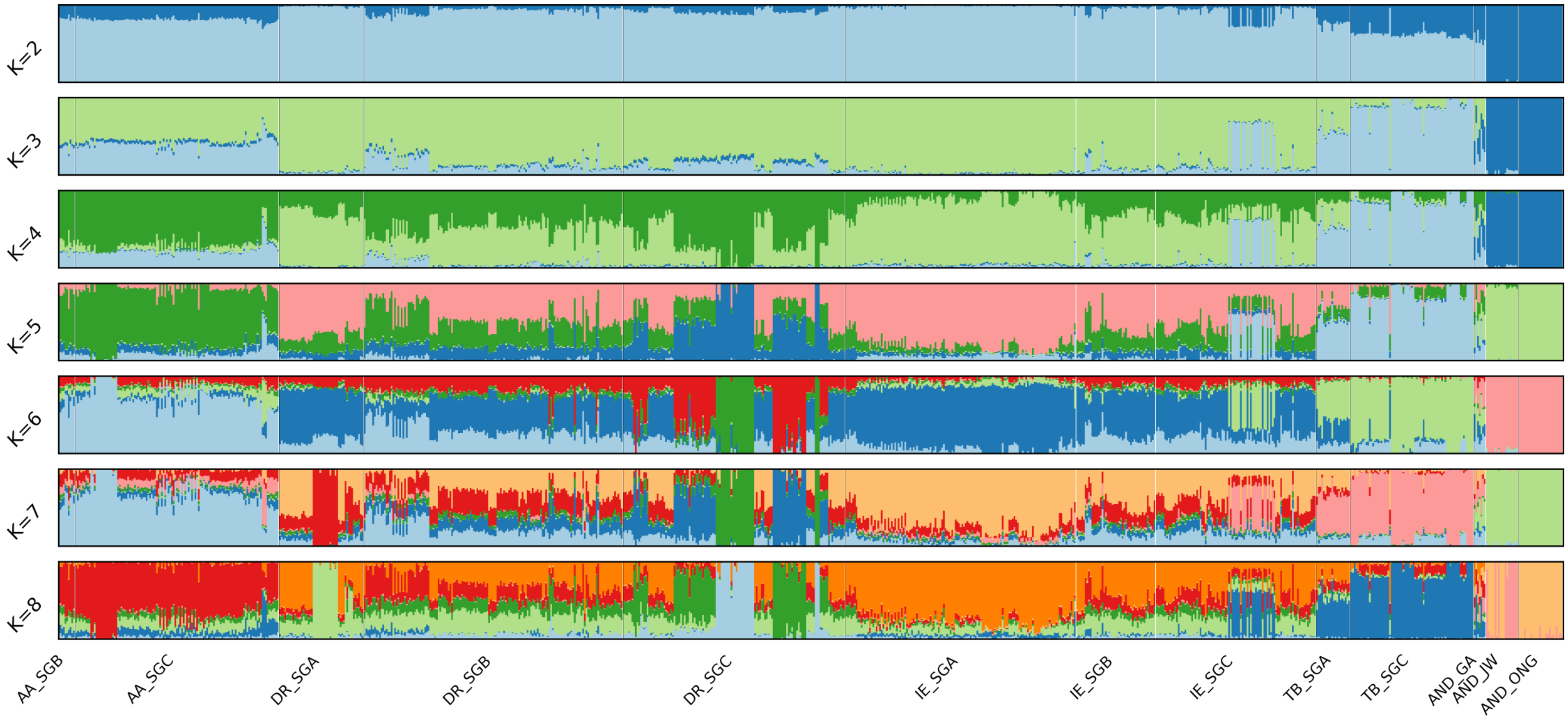


**Fig S3A**: An ADMIXTURE plot (for values of K between two and eight) of the mainland Indian data set (891 individuals; 47,283 SNPs) clearly shows the five main components related to language groups (Dravidian, Indo-European, Tibeto-Burman, Andamanese and Austro-Asiatic); see, for example, the plot for K equal to five or six. The plot also shows the divergence of the Dravidian SGC (DR_SGC) and the Andaman samples from rest of DR speakers and mainland India, respectively.

#### Fig S3B: Admixture ancestry of Indian populations

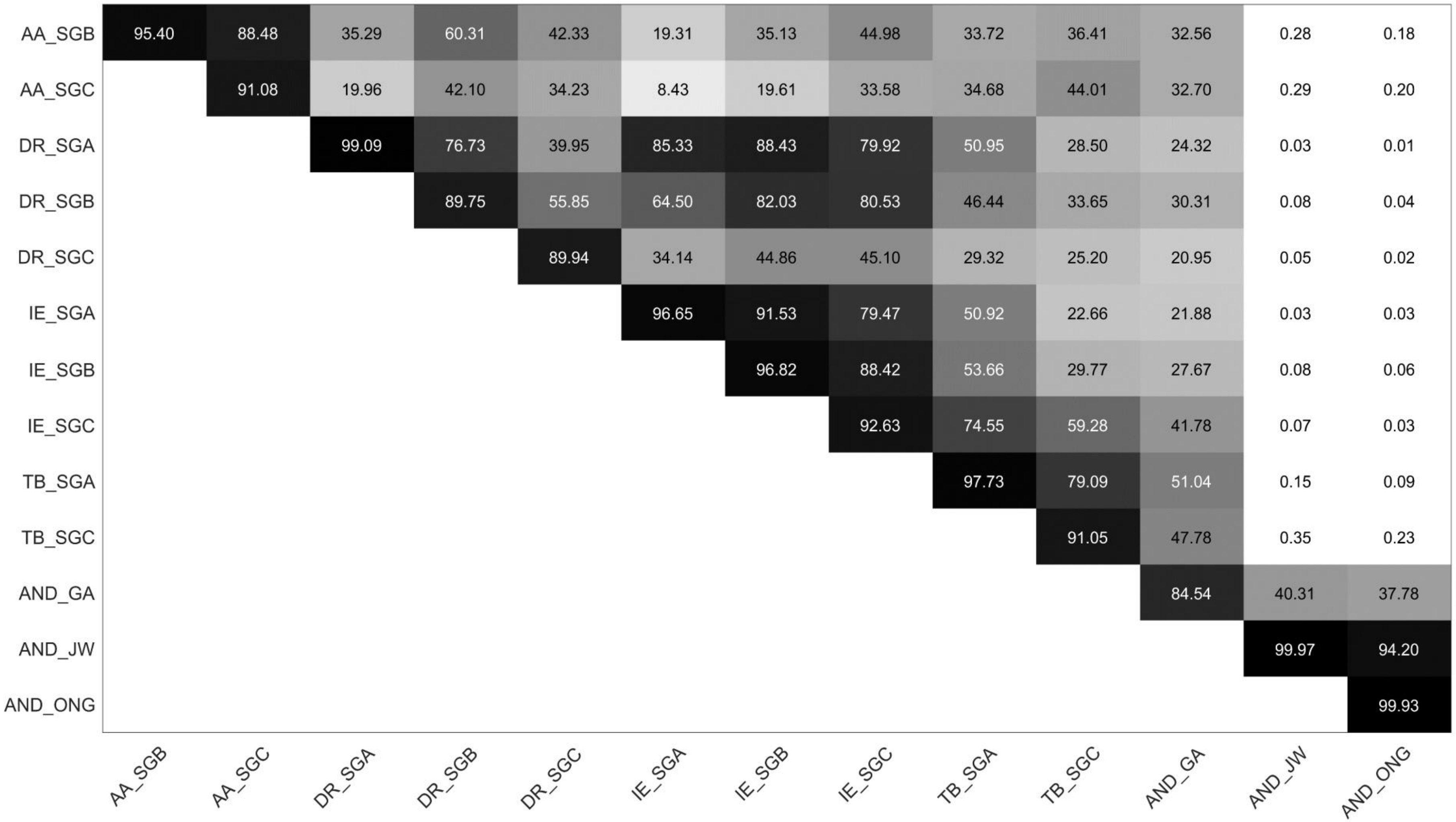


**Fig S3B**: Meta-analysis of the results of the ADMIXTURE plot (see Methods for details) to visually and numerically quantify the amount of shared ancestry (as revealed by ADMIXTURE) between any pair of populations. Darker colors indicate larger amounts of shared ancestry; we observe a higher amount of shared ancestry between the Indo-European and Dravidian populations, across all social groups, indicating the existence of significant admixture between the two linguistic groups. The isolation of the Dravidian SGC samples is primarily due to the isolation of hill SGCs (such as Irula, Kadar, Paniyas, etc.). Greater Andamanese (AND_GA) shares more ancestry with mainland Indian populations than other Andamanese groups Jarawa (AND_JW) and Onge (AND_ONG).

**Fig S4: Stratification in Indian populations**

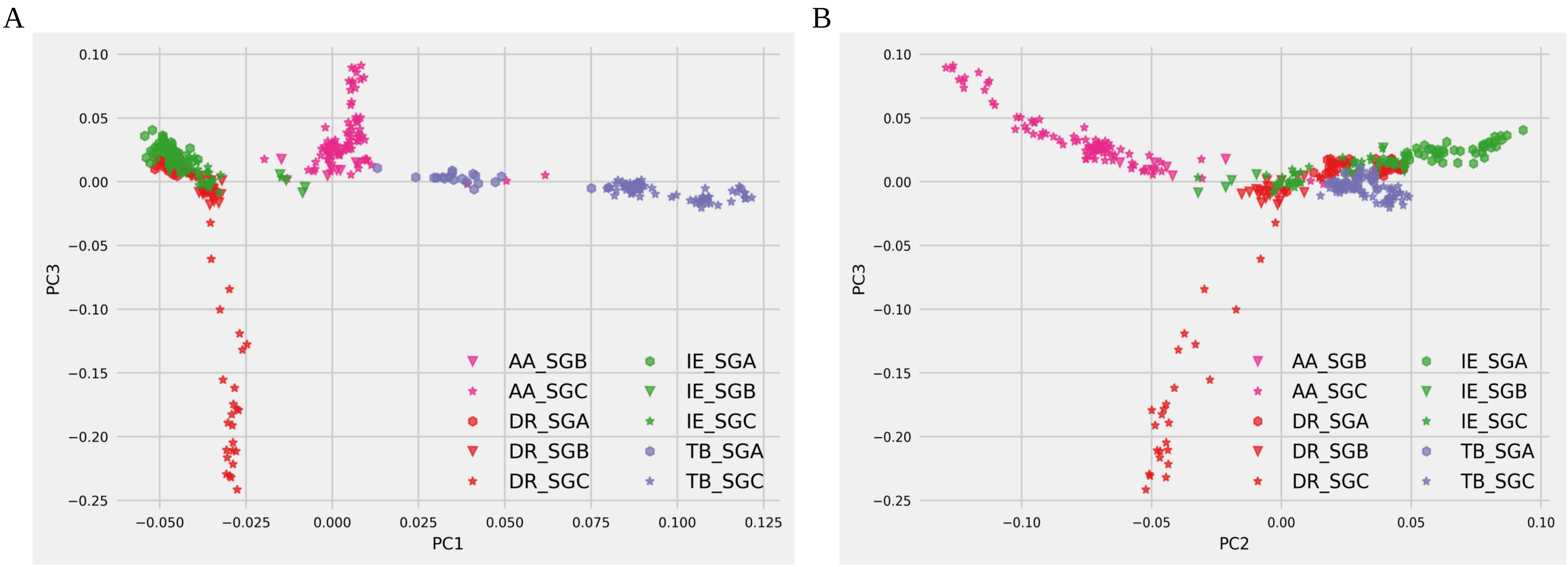


**Fig S4:** (A) First and third PCs reveal clusters stratified by sociolinguistic groups in the normalized data set of 368 individuals (33 populations). SGCs from different language groups diverge (IE_SGC is closer to other IE speakers) TB_SGC forms a cluster with TB_SGA being closer to IE and AA speakers. AA speakers form a cluster of their own.

(B) Second and third PCs extracted from the normalized data set reveal clear clusters by the sociolinguistic groups. DR_SGC shows divergence from fellow DR speakers (SGA and SGB) who tightly cluster with IE_SGB and IE_SGC. IE_SGA forms one end of a cline with maximal variance along with AA_SGC forming the other end with AA_SGB and IE_SGB possibly mixing in Central India.

**Fig S5: LDA plots**

A

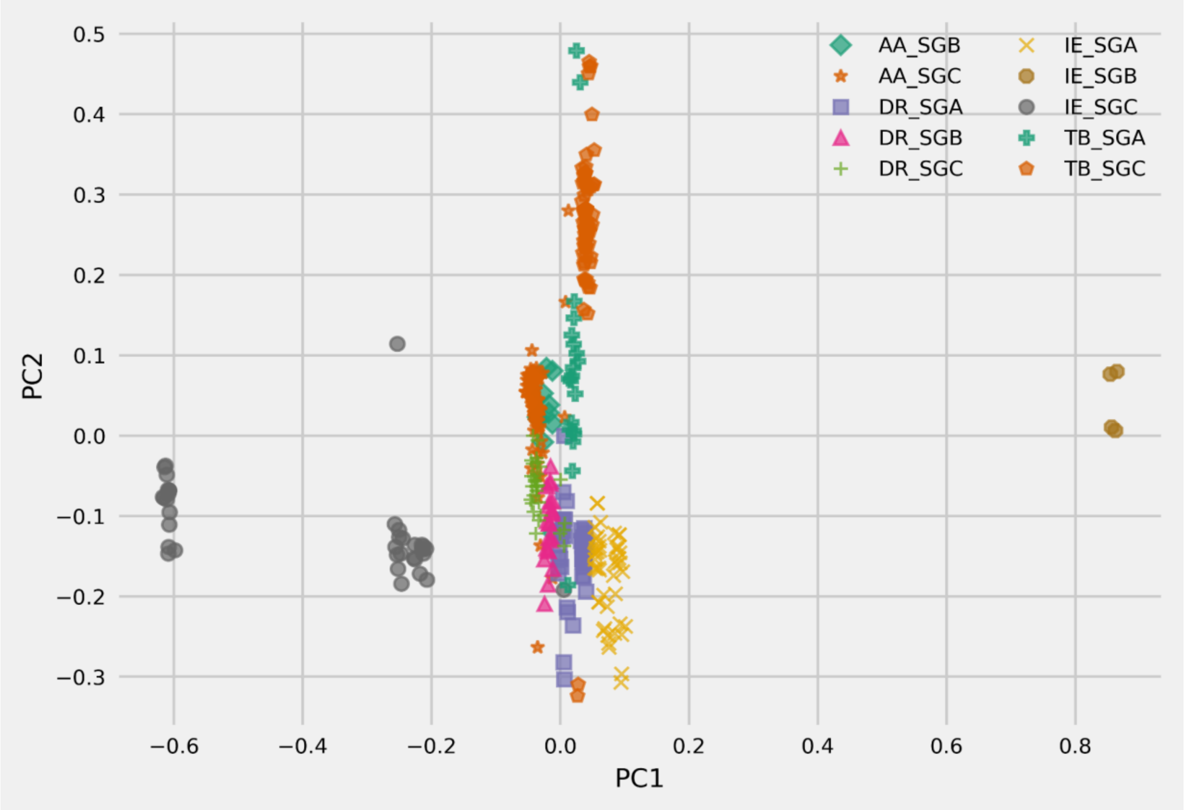


B

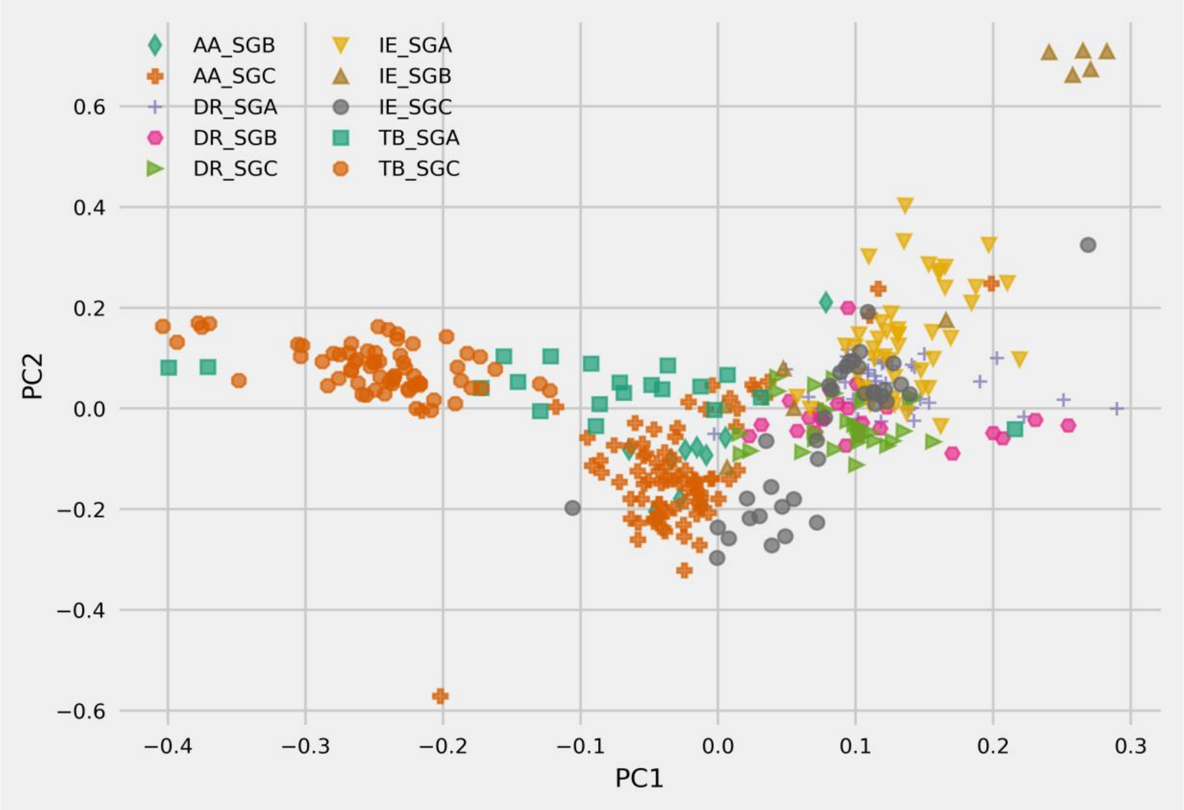


**Fig S5:** (A) Plotting the top two discriminants by language groups. Layers of stratification appear, from left to right. Although the LDA was performed by language groups, we see a two-layer stratification, first by castes and then by languages. The IE_SGA form a separate cline, followed by DR_SGA; then, the IE and DR SGBs follow. Then some DR and AA tribal populations cluster together, followed by a separate cluster of IE tribal populations.

(B) Plotting the top two discriminants by geographic regions. Layers of stratification appear from left to right. TB speakers occupy the left as IE speakers occupy the right side of the plot mirroring the east-west expanse of the map of India. However, the north-south variation does not appear as clearly as the east-west. This is perhaps confounded by the endogamy practiced by IE and DR populations.

**Fig S6: Statistical Significance of COGG**

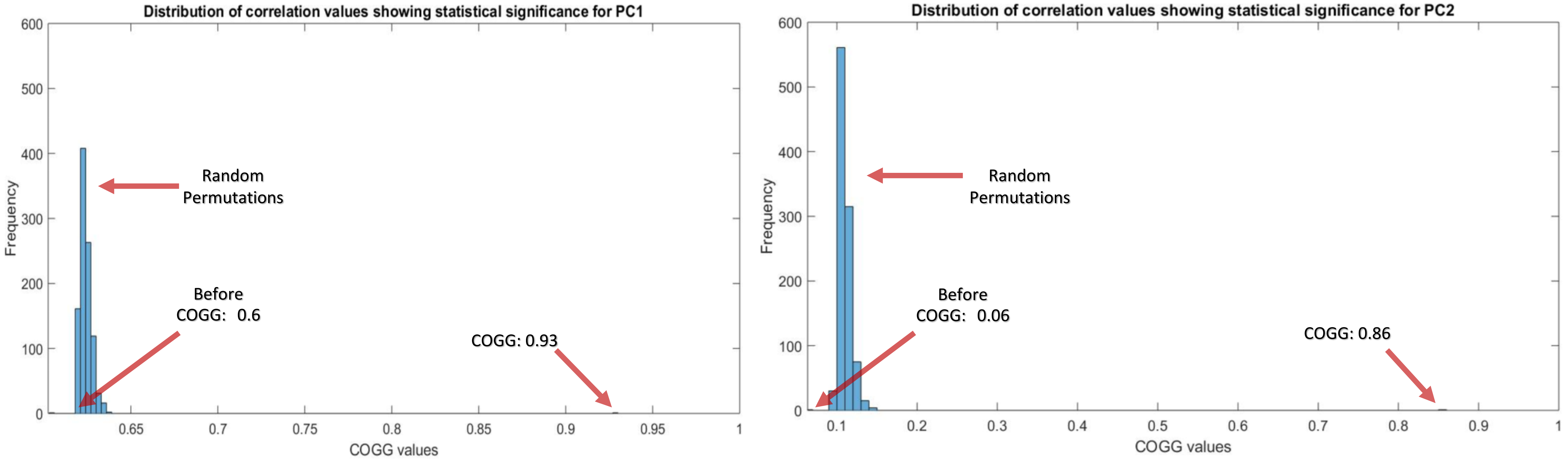


**Fig S6:** Statistical significance of the COGG output (using random permutations of the features) Clearly, COGG is statistically significant for both the first and the second principal components.

**Fig S7: COGG-CCA**

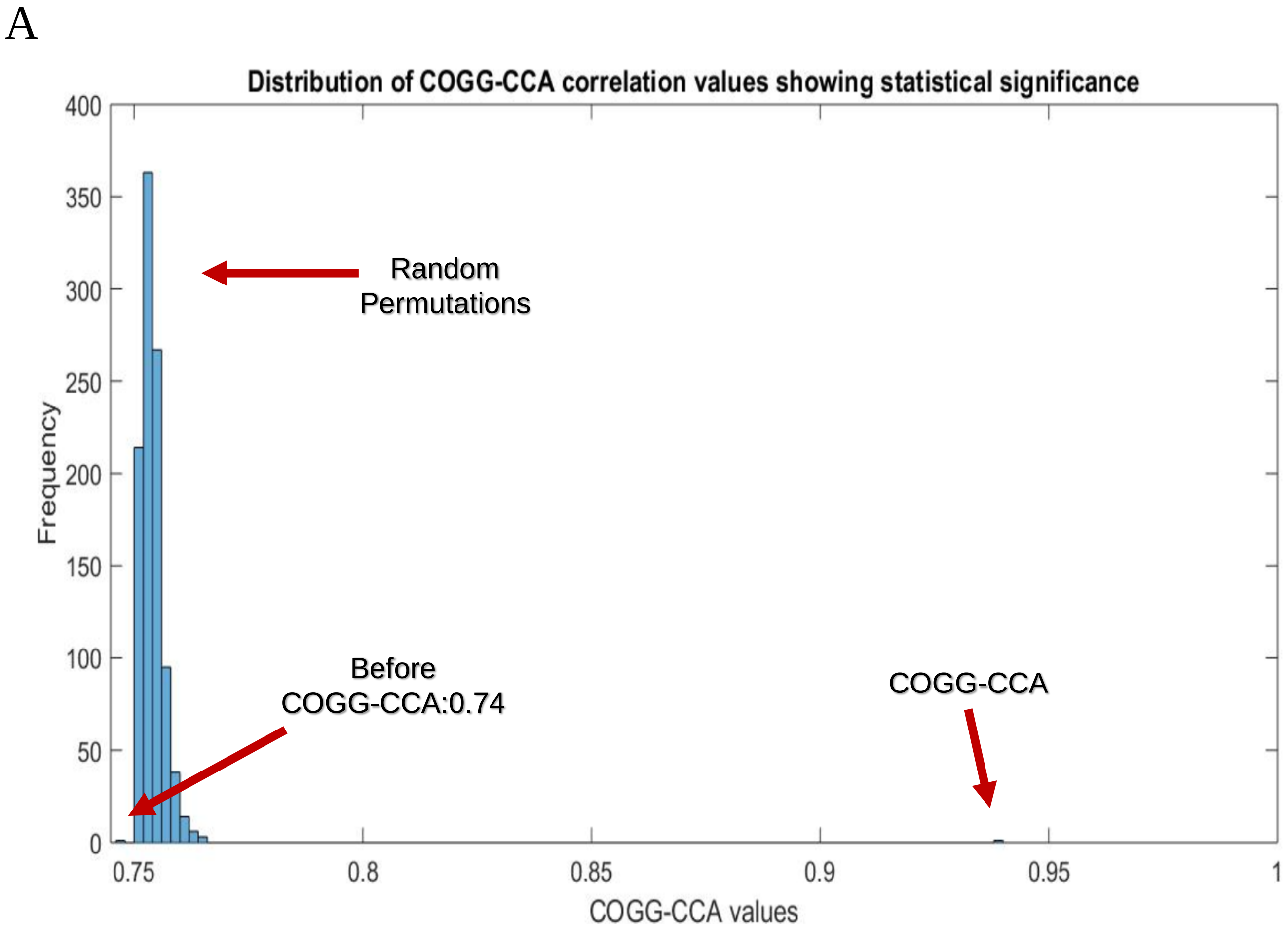


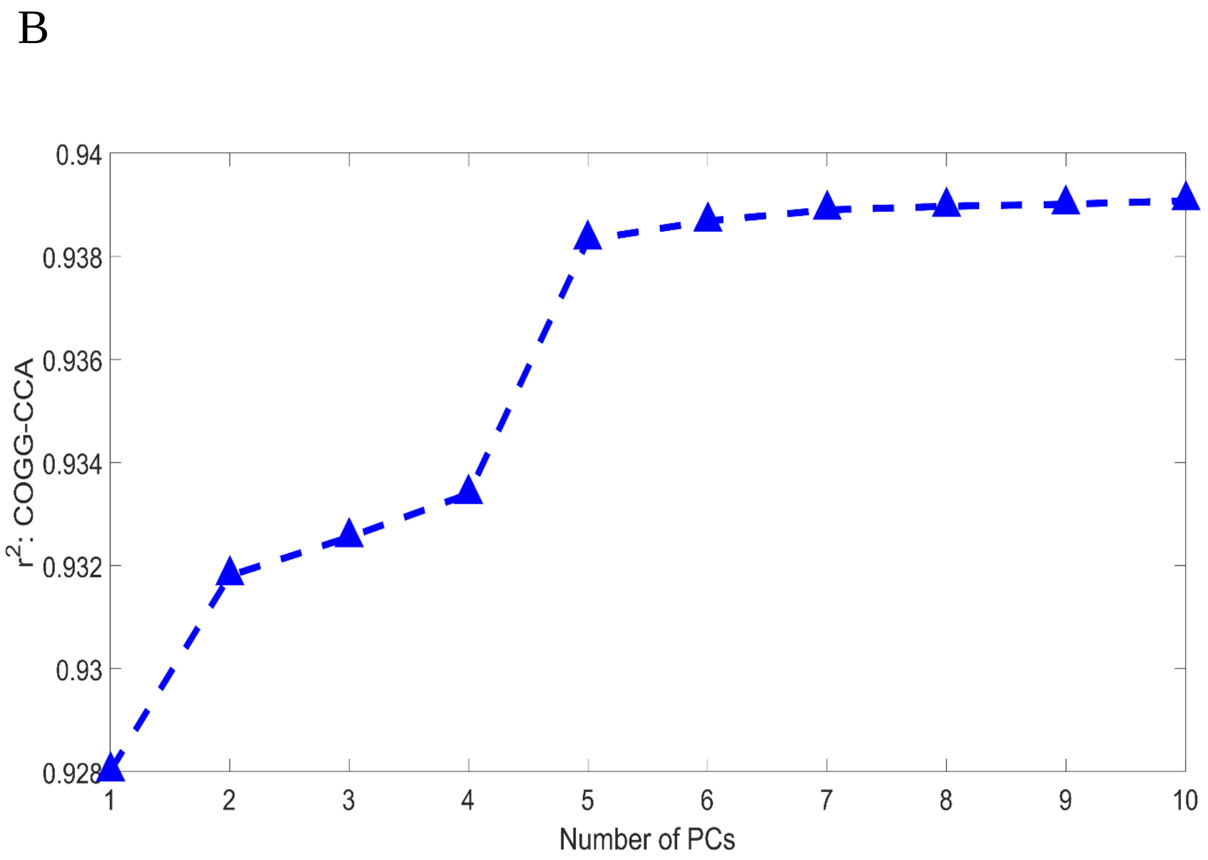


**Fig S7a**: COGG-CCA, when run with top 8 PCs, shows statistical significance with $r^2 = 0.94$ when compared against random permutations of the variables with average $r^2 = 0.75$.

**Fig S7b**: Varying number of PCs to perform COGG-CCA results in the maximum $r^2$ when top 6 to 8 PCs are used.

#### Fig S8: Shared Ancestry between IE and DR

| | Bhil_IE_SGC | Chamar_IE_SGC | Kanjars_IE_SGC | Lambadi_IE_SGC | Tharu_IE_SGC | Dharkar_IE_SGB | Dusadh_IE_SGB | Kurmi_IE_SGB | Lodi_IE_SGB | Meghawal_IE_SGB | Sahariya_IE_SGB | BrahminGJR_IE_SGA | Brahmin_IE_SGAs | BrahminUP_IE_SGA | BrahminUTR_IE_SGA | BrahminWB_IE_SGA | Kashmiri_Pandit_IE_SGA | Kshatriya_IE_SGA | Maratha_IE_SGA | Srivastava_IE_SGA |
|---|---|---|---|---|---|---|---|---|---|---|---|---|---|---|---|---|---|---|---|---|
| Adi-Dravider_DR_SGC | 94.052 | 95.133 | 68.246 | 78.055 | 2.460 | 66.806 | 92.518 | 71.172 | 85.063 | 64.252 | 82.964 | 48.103 | 38.757 | 53.207 | 38.275 | 59.364 | 33.025 | 48.917 | 94.616 | 66.857 |
| Hakkipikki_DR_SGC | 92.340 | 93.634 | 67.086 | 75.696 | 3.825 | 65.774 | 90.274 | 69.794 | 83.152 | 62.902 | 85.625 | 47.208 | 38.452 | 51.948 | 40.769 | 58.571 | 32.869 | 48.125 | 93.120 | 64.989 |
| Hallaki_DR_SGC | 92.855 | 92.308 | 83.384 | 88.546 | 1.918 | 82.415 | 92.311 | 85.048 | 91.371 | 81.074 | 68.883 | 69.306 | 61.140 | 73.379 | 48.079 | 77.614 | 55.664 | 69.835 | 92.716 | 82.615 |
| Irula_DR_SGC | 12.980 | 12.654 | 5.301 | 6.123 | 0.115 | 5.836 | 9.661 | 5.280 | 8.103 | 3.867 | 25.724 | 2.553 | 1.821 | 2.711 | 1.490 | 3.820 | 1.292 | 2.931 | 15.098 | 4.159 |
| Kadar_DR_SGC | 23.444 | 23.136 | 8.861 | 11.074 | 0.596 | 9.272 | 18.383 | 9.205 | 15.159 | 6.688 | 40.419 | 3.303 | 1.827 | 3.927 | 2.648 | 5.914 | 0.939 | 3.753 | 25.774 | 7.050 |
| Kuruchiyan_DR_SGC | 98.799 | 98.313 | 87.134 | 93.993 | 1.544 | 85.974 | 98.642 | 89.326 | 96.776 | 84.631 | 71.989 | 70.735 | 60.921 | 75.514 | 48.760 | 80.397 | 54.639 | 71.278 | 98.780 | 86.613 |
| Malli_DR_SGC | 94.011 | 92.004 | 97.071 | 98.493 | 1.420 | 96.563 | 93.781 | 97.801 | 97.980 | 95.876 | 60.310 | 87.012 | 79.208 | 90.274 | 56.841 | 93.581 | 73.422 | 87.367 | 93.784 | 96.308 |
| Palliyar_DR_SGC | 31.262 | 31.385 | 11.777 | 14.886 | 1.784 | 12.084 | 25.199 | 12.404 | 20.570 | 8.947 | 53.533 | 4.003 | 1.983 | 4.938 | 4.642 | 7.799 | 0.797 | 4.551 | 33.790 | 9.363 |
| Paniya_DR_SGC | 4.169 | 3.430 | 2.139 | 1.879 | 0.054 | 2.143 | 2.494 | 1.896 | 3.670 | 1.806 | 3.882 | 1.330 | 1.424 | 1.681 | 1.058 | 1.605 | 0.991 | 1.388 | 3.602 | 1.840 |
| Chenchu_DR_SGB | 32.377 | 33.961 | 25.308 | 27.549 | 7.525 | 24.835 | 31.695 | 25.865 | 30.545 | 23.273 | 32.855 | 19.005 | 17.016 | 20.409 | 24.301 | 22.826 | 15.986 | 19.679 | 31.512 | 25.261 |
| Kallar_DR_SGB | 99.075 | 98.251 | 87.174 | 93.552 | 1.415 | 86.178 | 98.045 | 89.033 | 96.930 | 84.384 | 72.682 | 70.631 | 61.003 | 75.430 | 48.262 | 80.254 | 54.477 | 71.265 | 99.084 | 86.342 |
| Kamsali_DR_SGB | 94.224 | 95.086 | 74.047 | 83.234 | 2.827 | 72.583 | 93.834 | 76.615 | 88.796 | 70.915 | 76.037 | 55.537 | 46.115 | 60.924 | 43.322 | 66.159 | 40.367 | 56.202 | 93.785 | 73.826 |
| Kurumba_DR_SGB | 81.000 | 80.323 | 65.732 | 71.011 | 1.626 | 65.116 | 78.099 | 67.337 | 75.527 | 62.588 | 69.397 | 51.131 | 43.719 | 54.646 | 35.180 | 59.294 | 38.577 | 51.814 | 82.030 | 63.841 |
| Madiga_DR_SGB | 83.602 | 85.858 | 55.331 | 66.272 | 2.166 | 53.562 | 84.100 | 58.655 | 73.052 | 51.878 | 76.708 | 37.122 | 29.482 | 41.602 | 30.756 | 46.927 | 25.535 | 37.759 | 84.170 | 54.488 |
| Mala_DR_SGB | 83.794 | 86.114 | 54.137 | 65.797 | 1.668 | 52.404 | 84.364 | 57.610 | 72.550 | 50.555 | 76.927 | 35.552 | 27.696 | 40.088 | 28.792 | 45.474 | 23.779 | 36.236 | 84.558 | 53.436 |
| Malayan_DR_SGB | 27.051 | 26.612 | 10.456 | 12.730 | 0.667 | 11.055 | 20.805 | 10.694 | 17.632 | 7.738 | 46.204 | 3.934 | 2.211 | 4.620 | 3.182 | 7.027 | 1.061 | 4.557 | 29.598 | 8.239 |
| Narikkuravar_DR_SGB | 95.114 | 93.354 | 93.582 | 97.638 | 0.476 | 92.419 | 96.360 | 95.466 | 97.118 | 92.726 | 59.786 | 81.894 | 73.156 | 85.589 | 49.816 | 88.869 | 67.228 | 81.904 | 95.109 | 93.092 |
| Pallan_DR_SGB | 93.602 | 94.286 | 70.868 | 79.418 | 2.610 | 69.674 | 91.679 | 73.377 | 85.890 | 67.048 | 81.462 | 51.845 | 42.815 | 56.679 | 40.508 | 62.622 | 37.034 | 52.673 | 94.125 | 69.356 |
| Sakilli_DR_SGB | 95.133 | 95.476 | 73.323 | 81.827 | 1.977 | 72.163 | 93.526 | 75.874 | 87.660 | 69.624 | 80.622 | 54.344 | 44.990 | 59.123 | 40.183 | 65.037 | 38.928 | 55.122 | 95.999 | 71.626 |
| TamilNadu_SC_DR_SGB | 97.247 | 95.446 | 92.039 | 96.232 | 0.724 | 91.374 | 96.345 | 93.424 | 97.476 | 89.858 | 66.469 | 78.414 | 69.402 | 82.314 | 48.996 | 86.378 | 63.034 | 78.928 | 97.595 | 90.972 |
| Iyer_DR_SGA | 84.926 | 81.920 | 98.714 | 96.830 | 0.621 | 98.709 | 85.302 | 98.366 | 93.243 | 98.905 | 46.181 | 95.084 | 89.476 | 96.973 | 56.311 | 97.938 | 84.984 | 95.187 | 84.402 | 98.712 |
| Naidu_DR_SGA | 97.092 | 95.247 | 90.498 | 95.243 | 0.611 | 89.742 | 95.960 | 91.918 | 97.122 | 88.257 | 65.753 | 76.529 | 67.534 | 80.784 | 47.364 | 84.517 | 61.114 | 77.033 | 97.043 | 89.823 |
| TamilNadu_Brahmin_DR_SGA | 84.535 | 81.262 | 98.645 | 96.067 | 0.612 | 98.911 | 84.206 | 98.086 | 92.512 | 98.539 | 46.761 | 95.303 | 89.955 | 96.790 | 55.743 | 97.953 | 85.462 | 95.551 | 84.356 | 98.127 |
| Vysya_DR_SGA | 36.187 | 37.651 | 26.120 | 31.753 | 0.314 | 25.899 | 37.109 | 26.706 | 33.745 | 25.196 | 27.677 | 21.156 | 18.884 | 23.030 | 16.212 | 23.755 | 17.893 | 21.542 | 34.898 | 28.059 |

**Fig S8:** The shared ancestry matrix of relatedness between IE and DR speakers show that high relatedness with some divergent groups, following from the PC plot in Fig S5a. The DR_SGA share very high ancestry with IE SGA and SGC, showing that there was high admixture and contact between these groups prior to endogamy.

**Fig S9: Shared ancestry of Indian populations**

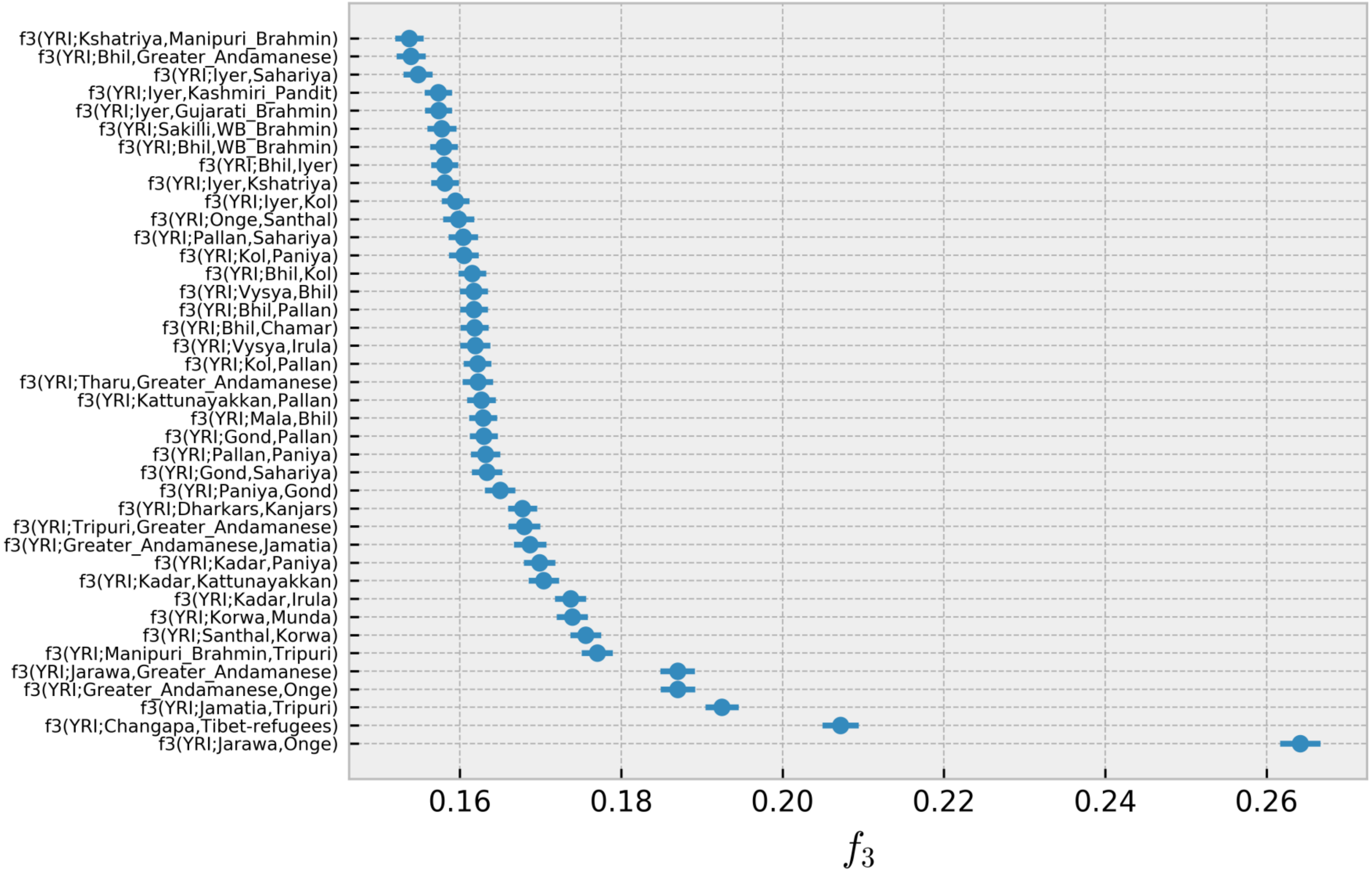


**Fig S9**:Most significant (Z-score higher than 85) outgroup $f_3$ statistics of the form f3(YRI;A,B) where YRI is the outgroup, A are the groups from Table S1 and B are all the pan-Indian populations in our data spanning across social groups and language families.

**Fig S10A: Network analysis of Eurasia in light of Indian populations**

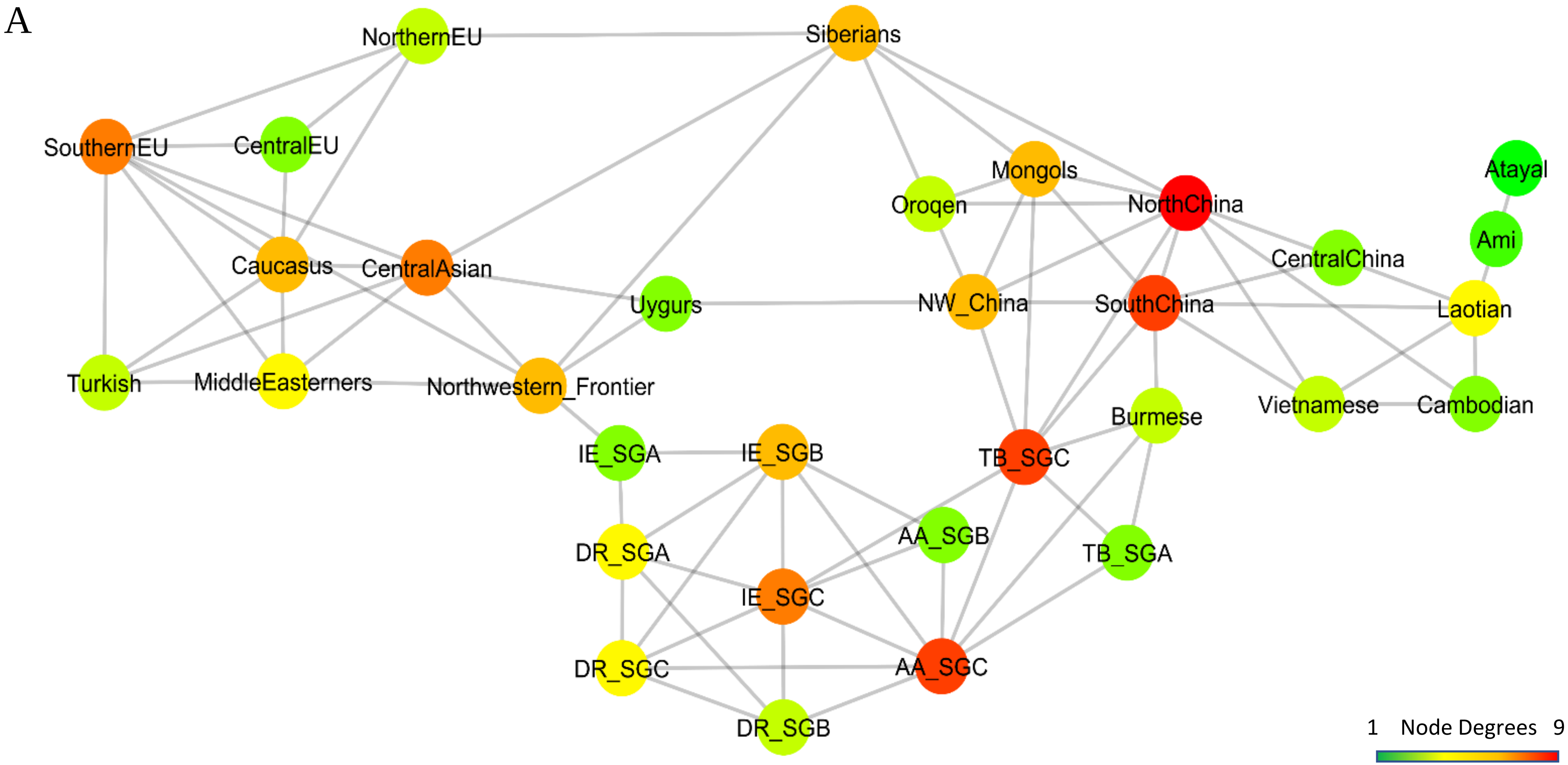


**Fig S10A**: Networks formed using the top five PCs (see Materials and Methods for the network formation algorithm) and five nearest neighbors showing three major paths leading to the two entry points of India.

#### Fig S10B: Shared ancestry of Eurasian populations

B

| | Northwestern_Frontier | CentralAsian | Turkish | MiddleEasterners | Caucasus | CentralEU | SouthernEU | NorthernEU | Mongols | Uygurs | Oroqen | Siberians |
|---|---|---|---|---|---|---|---|---|---|---|---|---|
| AA_SGB | 8.430 | 2.742 | 1.141 | 2.014 | 0.959 | 0.452 | 0.513 | 0.518 | 2.646 | 5.920 | 1.532 | 0.530 |
| AA_SGC | 3.523 | 0.939 | 0.318 | 0.563 | 0.221 | 0.227 | 0.189 | 0.291 | 2.452 | 3.562 | 1.560 | 0.528 |
| DR_SGB | 34.251 | 12.326 | 5.686 | 11.594 | 6.022 | 0.890 | 1.521 | 0.924 | 0.387 | 8.758 | 0.098 | 0.143 |
| DR_SGA | 43.508 | 21.496 | 13.815 | 21.819 | 14.741 | 4.651 | 6.561 | 4.215 | 0.393 | 12.074 | 0.061 | 0.141 |
| DR_SGC | 14.377 | 5.179 | 2.756 | 5.197 | 2.911 | 0.629 | 0.930 | 0.659 | 0.266 | 3.757 | 0.062 | 0.058 |
| IE_SGB | 50.482 | 27.747 | 18.132 | 26.354 | 18.505 | 6.883 | 9.482 | 6.247 | 2.364 | 18.444 | 1.168 | 0.754 |
| IE_SGA | 78.576 | 52.085 | 38.493 | 50.993 | 39.453 | 17.211 | 22.870 | 15.497 | 1.388 | 27.900 | 0.338 | 0.678 |
| IE_SGC | 46.453 | 21.930 | 13.046 | 21.102 | 13.521 | 4.168 | 5.870 | 3.883 | 0.628 | 13.638 | 0.151 | 0.167 |
| TB_SGA | 25.174 | 14.874 | 6.284 | 7.283 | 4.439 | 2.282 | 2.706 | 2.611 | 51.763 | 53.312 | 35.978 | 13.365 |
| TB_SGC | 6.423 | 4.062 | 0.751 | 0.578 | 0.260 | 0.133 | 0.155 | 0.286 | 65.076 | 44.139 | 48.949 | 19.362 |

**Fig S9B**: Meta-analysis of the ADMIXTURE output reveals that, overall, Indian populations share a great proportion of ancestry with the so-called Indian Northwestern Frontier populations, namely the SGC populations spanning Afghanistan and Pakistan. In concordance with previous studies we find higher degrees of shared ancestry in Central Asian populations with IE and DR SGA. In particular, IE SGA share large amounts of ancestry with other IE speaking populations (i.e., Europeans). However, IE, TB, and DR speakers also share considerable amounts of ancestry with the Uygurs. On the other hand, AA speakers, who have been suggested as the earliest settlers of India, appear more isolated.

| Population Name | # of Samples | State/Province | BroadRegion | Language | Caste | Latitude | Longitude | Dataset |
|---|---|---|---|---|---|---|---|---|
| **Adi-Dravider** | 5 | Tamil Nadu | South | Dravidian | Social Group B (SGB) | 12.11 | 79.053 | Moorjani et al. (2013) |
| **Aonaga** | 4 | Nagaland | NorthEast | Tibeto_Burmese | Unknown | 25.6667 | 94.133 | Reich et al. (2009) +Metspalu et al. (2011) |
| **Asur** | 2 | Jharkhand | Eastern States | Austro_Asiatic | Social Group C (SGC) | 23.76 | 86.42 | Chaubey et al. (2011) +Metspalu et al. (2011) |
| **Bhil** | 17 | Gujarat | NorthWest | Indo-European | SGC | 23.0333 | 72.667 | Moorjani et al. (2013) +Reich et al. (2009) |
| **Bhumij** | 5 | West Bengal | Eastern States | Austro_Asiatic | SGC | 21.806 | 87.114 | Moorjani et al. (2013) |
| **Bhunjia** | 1 | Odisha | Eastern States | Indo-European | SGC | 21.27 | 81.56 | Metspalu et al. (2011) |
| **Birhor** | 20 | Jharkhand | Eastern States | Austro_Asiatic | SGC | 23.991 | 84.816 | Basu et al. (2016) +Moorjani et al. (2013) |
| **Bonda** | 4 | Odisha | Eastern States | Austro_Asiatic | SGC | 18.4 | 81.88 | Metspalu et al. (2011) |
| **Brahmin** | 15 | Uttar Pradesh | North | Indo-European | Social Group A (SGA) | 25.75 | 82.683 | Moorjani et al. (2013) |
| **Chamar** | 10 | Bihar | Eastern States | Indo-European | SGC | 25.37 | 83.04 | Metspalu et al. (2011) |
| **Changapa** | 5 | Ladakh | North | Tibeto_Burmese | SGC | 34.02 | 79.004 | Moorjani et al. (2013) |
| **Chenchus** | 10 | Andhra Pradesh | South | Dravidian | SGC | 18 | 79.59 | Metspalu et al. (2011) + Reich et al. (2009) |
| **Dhurwa** | 1 | Bihar | Eastern States | Dravidian | SGC | 18.78 | 82.68 | Metspalu et al. (2011) |
| **Dharkars** | 12 | Uttar Pradesh | North | Indo-European | SGC | 25.44 | 83.1 | Metspalu et al. (2011) |
| **Dusadh** | 10 | Uttar Pradesh | North | Indo-European | SGB | 25.44 | 84.56 | Metspalu et al. (2011) |
| **Gadaba** | 1 | Andhra Pradesh | South | Austro_Asiatic | SGC | 18.79 | 82.7 | Chaubey et al. (2011) |
| **Garo** | 4 | Assam | NorthEast | Tibeto_Burmese | Unknown | 26.17 | 90.62 | Metspalu et al. (2011) |
| **Gond** | 38 | Madhya Pradesh | Central | Dravidian | SGC | 22.1 | 82.16 | Reich et al. (2009) +Metspalu et al. (2011) +Basu et al. (2016) |
| **Gounder** | 5 | Tamil Nadu | South | Dravidian | SGA | 12.1 | 79.1 | Moorjani et al. (2013) |
| **Gujarati_Brahmin** | 20 | Gujarat | NorthWest | Indo-European | SGA | 22.29 | 70.94 | Basu et al. (2016) |
| **Hallaki** | 7 | Kannada | South | Dravidian | SGC | 13.9167 | 74.15 | Reich et al. (2009) |
| **Hakkipikki** | 4 | Kannada | South | Dravidian | SGC | 14.78 | 74.51 | Metspalu et al. (2011) |
| **Ho** | 28 | Jharkhand | Eastern States | Austro_Asiatic | SGC | 25.4 | 86.13 | Reich et al. (2009) +Metspalu et al. (2011) +Basu et al. (2016) |
| **Irula** | 25 | Tamil Nadu | South | Dravidian | SGC | 11.58 | 76.609 | Basu et al. (2016) +Moorjani et al. (2013) |
| **Iyer** | 20 | Tamil Nadu | South | Dravidian | SGA | 13.1 | 80.2 | Basu et al. (2016) |
| **Jamatia** | 18 | Tripura | NorthEast | Tibeto_Burmese | SGC | 23.84 | 92.17 | Basu et al. (2016) |
| **Juang** | 2 | Odisha | Eastern States | Austro_Asiatic | SGC | 21.49 | 83.98 | Metspalu et al. (2011) |
| **Kadar** | 20 | Kerala | South | Dravidian | SGC | 9.96 | 77.16 | Basu et al. (2016) |
| **Kallar** | 5 | Tamil Nadu | South | Dravidian | SGC | 10.99 | 78.22 | Metspalu et al. (2011) +Moorjani et al. (2013) |
| **Kamsali** | 4 | Andhra Pradesh | South | Dravidian | SGB | 15.49 | 78.29 | Reich et al. (2009) |
| **Kanjars** | 8 | Rajasthan | North | Indo-European | SGC | 26.45 | 80.32 | Metspalu et al. (2011) |
| **Kashmiri_Pandit** | 20 | Kashmir | North | Indo-European | SGA | 34.22 | 75.5 | Reich et al. (2009) +Moorjani et al. (2013) |
| **Kattunayakkan** | 5 | Andhra Pradesh | South | Dravidian | SGC | 9.55 | 76.8 | Moorjani et al. (2013) |
| **Kharia** | 8 | Bihar | Eastern States | Austro_Asiatic | SGC | 21.89 | 83.36 | Metspalu et al. (2011) +Reich et al. (2009) |
| **Khasi** | 3 | Meghalaya | NorthEast | Austro_Asiatic | SGC | 24.87 | 90.72 | Metspalu et al. (2011) |
| **Khatri** | 19 | Punjab | North | Indo-European | SGA | 30.52 | 76.76 | Basu et al. (2016) |
| **Kol** | 17 | Uttar Pradesh | North | Indo-European | SGB | 25.15 | 82.58 | Metspalu et al. (2011) |
| **Korku** | 4 | Madhya Pradesh | Central | Austro_Asiatic | SGB | 22.711 | 75.88 | Moorjani et al. (2013) |

| | | | | | | | | |
|---|---|---|---|---|---|---|---|---|
| **Korwa** | 18 | Jharkhand | Eastern States | Austro_Asiatic | SGC | 22.39 | 82.79 | Basu et al. (2016) |
| **Kshatriya** | 27 | Uttar Pradesh | North | Indo-European | SGA | 25.45 | 82.41 | Moorjani et al. (2013) +Metspalu et al. (2011) |
| **Kurmi** | 1 | West Bengal | East | Indo-European | SGB | 22.85 | 88.3 | Metspalu et al. (2011) |
| **Kuruchiyan** | 5 | Kerala | South | Dravidian | SGC | 11.73 | 76.41 | Moorjani et al. (2013) |
| **Kurumba** | 13 | Tamil Nadu | South | Dravidian | SGC | 10.54 | 76.27 | Reich et al. (2009) +Metspalu et al. (2011) |
| **Lambadi** | 1 | Madhya Pradesh | Central | Dravidian | SGC | 17.45 | 78.5 | Metspalu et al. (2011) |
| **Lodi** | 5 | Uttar Pradesh | North | Indo-European | SGB | 26.45 | 83.24 | Reich et al. (2009) |
| **Madiga** | 19 | Andhra Pradesh | South | Dravidian | SGB | 17.58 | 79.35 | Moorjani et al. (2013) +Reich et al. (2009) |
| **Mala** | 18 | Andhra Pradesh | South | Dravidian | SGB | 17.22 | 78.29 | Moorjani et al. (2013) +Reich et al. (2009) |
| **Malayan** | 2 | Tamil Nadu | South | Dravidian | SGC | 9.58 | 76.51 | Metspalu et al. (2011) |
| **Malai_Kuravar** | 5 | Tamil Nadu | South | Dravidian | SGB | 13.84 | 80.22 | Moorjani et al. (2013) |
| **Malli** | 5 | Andhra Pradesh | South | Dravidian | SGB | 10.55 | 72.63 | Moorjani et al. (2013) |
| **Manipuri_Brahmin** | 20 | Manipur | NorthEast | Tibeto_Burmese | SGA | 24.812 | 93.94 | Basu et al. (2016) |
| **Mawasi** | 1 | Madhya Pradesh | Central | Austro_Asiatic | SGB | 23.15 | 77.42 | Basu et al. (2016) |
| **Maratha** | 7 | Maharashtra | West | Indo-European | SGA | 18.5 | 73.7 | Basu et al. (2016) |
| **Meghawal** | 6 | Gujarat | NorthWest | Indo-European | SGB | 26.18 | 73.04 | Reich et al. (2009) +Metspalu et al. (2011) |
| **Meena** | 1 | Rajasthan | NorthWest | Indo-European | SGC | 28.29 | 74.98 | Metspalu et al. (2011) |
| **Minicoy** | 5 | Lakshwadeep | SouthWest | Indo-European | SGB | 8.28 | 73.06 | Moorjani et al. (2013) |
| **Munda** | 5 | Jharkhand | Eastern States | Austro_Asiatic | SGB | 21.6 | 83.76 | Moorjani et al. (2013) |
| **Naga** | 4 | Nagaland | NorthEast | Tibeto_Burmese | SGC | 25.67 | 94.11 | Metspalu et al. (2011) |
| **Naidu** | 4 | Andhra Pradesh | South | Dravidian | SGA | 13.13 | 79.06 | Reich et al. (2009) |
| **Narikkuravar** | 5 | Tamil Nadu | South | Dravidian | SGC | 13.17 | 79.4 | Moorjani et al. (2013) |
| **Nysha** | 4 | Arunachal Pradesh | NorthEast | Tibeto_Burmese | SGC | 26.55 | 92.4 | Reich et al. (2009) |
| **Pallan** | 20 | Tamil Nadu | South | Dravidian | SGA | 9.92 | 78.12 | Basu et al. (2016) |
| **Palliyar** | 5 | Tamil Nadu | South | Dravidian | SGC | 10.89 | 76.84 | Moorjani et al. (2013) |
| **Pulliyar** | 5 | Tamil Nadu | South | Dravidian | SGB | 11.02 | 76.98 | Metspalu et al. (2011) |
| **Piramalai_Kallars** | 8 | Tamil Nadu | South | Dravidian | SGC | 10.99 | 78.22 | Metspalu et al. (2011) |
| **Paniyas** | 27 | Kerala | South | Dravidian | SGC | 9.5 | 76.8 | Moorjani et al. (2013) + Metspalu et al. (2011) + Basu et al. (2016) |
| **Sahariya** | 4 | Madhya Pradesh | Central | Indo-European | SGB | 25.28 | 81.54 | Reich et al. (2009) |
| **Sakilli** | 4 | Tamil Nadu | South | Dravidian | SGB | 9.86 | 76.97 | Metspalu et al. (2011) |
| **Santhal** | 28 | Jharkhand | Central+East | Austro_Asiatic | SGC | 24.3 | 87.3 | Metspalu et al. (2011) +Reich et al. (2009) +Basu et al. (2016) |
| **Satnami** | 4 | Madhya Pradesh | Central | Indo-European | SGB | 20.29 | 85.58 | Reich et al. (2009) |
| **Savara** | 2 | Odisha | Central+East | Austro_Asiatic | SGB | 18.8 | 82.7 | Metspalu et al. (2011) |
| **Sherpa** | 5 | Nepal | NorthEast | Tibeto_Burmese | SGC | 29.2 | 83.4 | Moorjani et al. (2013) |
| **Srivastava** | 2 | Uttar Pradesh | North | Indo-European | SGA | 25.1 | 82.37 | Reich et al. (2009) |
| **Subba** | 5 | Sikkim | NorthEast | Tibeto_Burmese | SGC | 27.34 | 88.6 | Moorjani et al. (2013) |
| **Tharu** | 31 | Nepal | North | Indo-European | SGC | 29.23 | 79.3 | Reich et al. (2009) +Basu et al. (2016) + Metspalu et al. (2011) |
| **Tibet-refugees** | 5 | Tibet | North | Tibeto_Burmese | SGC | 29.625 | 91.17 | Moorjani et al. (2013) |
| **Tripuri** | 19 | Tripura | NorthEast | Tibeto_Burmese | SGC | 23.81 | 91.2 | Basu et al. (2016) |
| **Vaish** | 4 | Uttar Pradesh | North | Indo-European | SGA | 25.46 | 82.44 | Reich et al. (2009) |
| **Vedda** | 4 | Sri Lanka | SriLanka | Indo-European | SGC | 6.44 | 80.5 | Moorjani et al. (2013) |
| **Velamas** | 14 | Andhra Pradesh | South | Dravidian | SGA | 17.05 | 79.27 | Reich et al. (2009) + Metspalu et al. (2011) |
| **Vysya** | 20 | Tamil Nadu | South | Dravidian | SGA | 14.41 | 77.39 | Reich et al. (2009) +Moorjani et al. (2013) |
| **WB_Brahmin** | 18 | West Bengal | East | Indo-European | SGA | 22.55 | 88.37 | Basu et al. (2016) |

| | | | | | | | | |
|---|---|---|---|---|---|---|---|---|
| **UttarPradesh_SC** | 5 | Uttar Pradesh | North | Indo-European | SGB | 25.42 | 83.1 | Metspalu et al. (2011) |
| **TamilNadu_SC** | 2 | Tamil Nadu | South | Dravidian | SGB | 13.05 | 80.18 | Metspalu et al. (2011) |
| **UttarPradesh_Brahmins** | 8 | Uttar Pradesh | North | Indo-European | SGA | 26.06 | 83.18 | Metspalu et al. (2011) |
| **Uttaranchal_Brahmins** | 1 | Uttar Pradesh | North | Indo-European | SGA | 29.6 | 79.65 | Metspalu et al. (2011) |
| **Jarawa** | 19 | Andaman | Andaman | Ongan | SGC | 11.7 | 92.6 | Basu et al. (2016) |
| **Onge** | 26 | Andaman | Andaman | Ongan | SGC | 11.7 | 92.6 | Basu et al. (2016) |
| **Great Andmanese** | 7 | Andaman | Andaman | Great Andamanese | SGC | 12.2 | 93 | Reich et al. (2009) |
| **TamilNadu_Brahmin** | 2 | Tamil Nadu | South | Dravidian | SGA | 12.49 | 78.42 | Metspalu et al. (2011) |

**Table S1A:** A detailed description of the Indian samples, including their place of origin, language, caste affiliations, and respective longitude/latitude. The last column references the publication describing the respective dataset (we use the first author's last name and year of publication as a shortcut to the relevant reference from our bibliography).

| Population Name | # of Samples | State/Province | BroadRegion | Language | Caste | Latitude | Longitude | Dataset |
|---|---|---|---|---|---|---|---|---|
| **Bhil** | 17 | Gujarat | NorthWest | Indo-European | SGC | 23.0333 | 72.6667 | Moorjani et al. (2013) [6]+Reich et al. (2009) |
| **Kanjars** | 8 | Rajasthan | North | Indo-European | SGC | 26.45 | 80.32 | Metspalu et al. (2011) |
| **Kashmiri_Pandit** | 20 | Kashmir | North | Indo-European | SGA | 34.22 | 75.5 | Reich et al. (2009) +Moorjani et al. (2013) [6] |
| **Chamar** | 10 | Bihar | Eastern States | Indo-European | SGC | 25.37 | 83.04 | Metspalu et al. (2011) |
| **Kshatriya** | 27 | Uttar Pradesh | North | Indo-European | SGA | 25.45 | 82.41 | Moorjani et al. (2013) [6]+Metspalu et al. (2011) |
| **Meghawal** | 6 | Gujarat | NorthWest | Indo-European | SGB | 26.18 | 73.04 | Reich et al. (2009) +Metspalu et al. (2011) |
| **Tharus** | 2 | Nepal | Central | Indo-European | SGC | 27.12 | 83.45 | Metspalu et al. (2011) |
| **Sahariya** | 4 | Madhya Pradesh | Central | Indo-European | SGB | 25.28 | 81.54 | Reich et al. (2009) |
| **Sherpa** | 5 | Nepal | NorthEast | Tibeto_Burmese | SGC | 29.2 | 83.4 | Moorjani et al. (2013) [6] |
| **Changapa** | 5 | Ladakh | North | Tibeto_Burmese | SGC | 34.02 | 79.004 | Moorjani et al. (2013) [6] |
| **Nysha** | 4 | Arunachal Pradesh | NorthEast | Tibeto_Burmese | SGC | 26.55 | 92.4 | Reich et al. (2009) |
| **Jamatia** | 18 | Tripura | NorthEast | Tibeto_Burmese | SGC | 23.84 | 92.17 | Basu et al. (2016) |
| **Aonaga** | 4 | Nagaland | NorthEast | Tibeto_Burmese | SGC | 25.6667 | 94.1333 | Reich et al. (2009) +Metspalu et al. (2011) |
| **Naga** | 4 | Nagaland | NorthEast | Tibeto_Burmese | SGC | 25.67 | 94.11 | Metspalu et al. (2011) |
| **Tripuri** | 19 | Tripura | NorthEast | Tibeto_Burmese | SGC | 23.81 | 91.2 | Basu et al. (2016) |
| **Manipuri_Brahmin** | 20 | Manipur | NorthEast | Tibeto_Burmese | SGA | 24.812 | 93.94 | Basu et al. (2016) |
| **Tibet-refugees** | 5 | Tibet | North | Tibeto_Burmese | SGC | 29.625 | 91.17 | Moorjani et al. (2013) [6] |
| **Subba** | 5 | Sikkim | NorthEast | Tibeto_Burmese | SGC | 27.34 | 88.6 | Moorjani et al. (2013) [6] |
| **Khasi** | 3 | Meghalaya | NorthEast | Austro_Asiatic | SGC | 24.87 | 90.72 | Metspalu et al. (2011) |
| **Bhumij** | 5 | West Bengal | Eastern States | Austro_Asiatic | SGC | 21.806 | 87.114 | Moorjani et al. (2013) [6] |
| **Birhor** | 20 | Jharkhand | Eastern States | Austro_Asiatic | SGC | 23.991 | 84.816 | Basu et al. (2016) +Moorjani et al. (2013) [6] |
| **Munda** | 5 | Jharkhand | Eastern States | Austro_Asiatic | SGB | 21.6 | 83.76 | Moorjani et al. (2013) [6] |
| **Mawasi** | 1 | Madhya Pradesh | Central | Austro_Asiatic | SGB | 23.15 | 77.42 | Basu et al. (2016) |
| **Santhal** | 28 | Jharkhand | Central+East | Austro_Asiatic | SGC | 24.3 | 87.3 | Metspalu et al. (2011) +Reich et al. (2009) +Basu et al. (2016) |
| **Kharia** | 8 | Bihar | Eastern States | Austro_Asiatic | SGC | 21.89 | 83.36 | Metspalu et al. (2011) +Reich et al. (2009) |
| **Korku** | 4 | Madhya Pradesh | Central | Austro_Asiatic | SGB | 22.711 | 75.88 | Moorjani et al. (2013) [6] |
| **Korwa** | 18 | Jharkhand | Eastern States | Austro_Asiatic | SGC | 22.39 | 82.79 | Basu et al. (2016) |
| **Sakilli** | 4 | Tamil Nadu | South | Dravidian | SGB | 9.86 | 76.97 | Metspalu et al. (2011) |
| **Irula** | 25 | Tamil Nadu | South | Dravidian | SGC | 11.58 | 76.609 | Basu et al. (2016)+Moorjani et al. (2013) [6] |
| **Kuruchiyan** | 5 | Kerala | South | Dravidian | SGC | 11.73 | 76.41 | Moorjani et al. (2013) [6] |
| **Madiga** | 19 | Andhra Pradesh | South | Dravidian | SGB | 17.58 | 79.35 | Moorjani et al. (2013) [6]+Reich et al. (2009) |
| **Vysya** | 20 | Tamil Nadu | South | Dravidian | SGA | 14.41 | 77.39 | Reich et al. (2009) +Moorjani et al. (2013) [6] |
| **Iyer** | 20 | Tamil Nadu | South | Dravidian | SGA | 13.1 | 80.2 | Basu et al. (2016) |

**Table S1B:** Normalized subset of samples in India created after carefully selecting populations from Table S1A to equally represent, region, caste and languages. For each population, we include their place of origin, language, caste affiliations, and respective longitude/latitude. The last column references the publication describing the respective dataset (we use the first author's last name and year of publication as a shortcut to the relevant reference).

| Population Name | # of samples | Region | Data Source |
|---|---|---|---|
| **Adygei** | 38 | Caucasus | Cann et al. (2002) +Rajeevan et al. (2003) |
| **Afghan** | 24 | NW_Frontier | Cann et al. (2002) + Di Cristofaro et al. (2013) |
| **Albania** | 30 | SouthernEU | Rajeevan et al. (2003) |
| **Ami** | 38 | SouthEast Asia | Rajeevan et al. (2003) |
| **Atayal** | 34 | SouthEast Asia | Rajeevan et al. (2003) |
| **Azeris** | 23 | CentralAsian | Yunusbayev et al. (2015) |
| **Bedouin** | 48 | MiddleEast | Cann et al. (2002) |
| **Brahui** | 25 | NW_Frontier | Cann et al. (2002) Di Cristofaro et al. (2013) |
| **Burmese** | 15 | Burmese | Chaubey et al. (2011) |
| **Burusho** | 25 | NW_Frontier | Cann et al. (2002) Di Cristofaro et al. (2013) |
| **Buryats** | 22 | Siberian | Yunusbayev et al. (2015) |
| **Cambodian** | 26 | SouthEast Asia | Cann et al. (2002) |
| **Chechens** | 20 | Caucasus | Yunusbayev et al. (2012) |
| **Druze** | 50 | MiddleEast | Cann et al. (2002) |
| **French** | 29 | CentralEU | Cann et al. (2002) |
| **Georgians** | 30 | Caucasus | Yunusbayev et al. (2012) |
| **Germans** | 13 | CentralEU | Yunusbayev et al. (2015) |
| **Greek** | 20 | SouthernEU | Behar et al. (2012) +d11 |
| **Hakka** | 37 | SouthChina | Rajeevan et al. (2003) |
| **Han** | 44 | NorthChina | Cann et al. (2002) |
| **Hazara** | 24 | NW_Frontier | Cann et al. (2002) + Di Cristofaro et al. (2013) |
| **Hezhen** | 9 | NorthChina | Cann et al. (2002) |
| **Iranians** | 20 | MiddleEast | Behar et al. (2012) |
| **Ishkashim** | 10 | NW_Frontier | Cann et al. (2002) + Di Cristofaro et al. (2013) |
| **Italian** | 37 | SouthernEU | Cann et al. (2002) + Behar et al. (2012) |
| **KHV** | 99 | SouthEast Asia | Behar et al. (2012) |
| **Kabardin** | 3 | Caucasus | Auton et al. (2015) [41] |
| **Kurds** | 6 | MiddleEast | Yunusbayev et al. (2012) |
| **Laotians** | 59 | SouthEast Asia | Rajeevan et al. (2003) |
| **Lebanese** | 8 | MiddleEast | Behar et al. (2012) |
| **Libya** | 17 | MiddleEast | Rajeevan et al. (2003) |
| **Mongolians** | 21 | Mongolia | Cann et al. (2002) |
| **Naxi** | 9 | SouthChina | Cann et al. (2002) |
| **Oroqen** | 10 | NorthChina | Cann et al. (2002) |
| **Romanians** | 32 | SouthernEU | Behar et al. (2012) |
| **Russians** | 83 | NorthernEU | Cann et al. (2002) +Rajeevan et al. (2003) +Yunusbayev et al. (2015) |
| **Selkups** | 20 | Siberian | Raghavan et al. (2014) |
| **She** | 10 | SouthChina | Cann et al. (2002) |
| **Swedish** | 18 | NorthernEU | Behar et al. (2012) |
| **Syrians** | 16 | MiddleEast | Behar et al. (2012) |
| **Tajiks** | 24 | CentraAsian | Yunusbayev et al. (2015) +Yunusbayev et al. (2012) |
| **Tu** | 10 | NW_China | Cann et al. (2002) |
| **Tujia** | 10 | CentralChina | Cann et al. (2002) |
| **Turkmens** | 23 | CentralAsian | Behar et al. (2012) +Yunusbayev et al. (2012) |
| **Turks** | 19 | MiddleEast | Yunusbayev et al. (2015) +Cann et al. (2002) |
| **Uyghur** | 11 | Uyghurs | Behar et al. (2012) |
| **Uzbeks** | 19 | CentralAsian | Cann et al. (2002) |
| **Xibo** | 9 | NW_China | Rajeevan et al. (2003) |
| **Yakuts** | 49 | Siberian | Behar et al. (2012) |
| **Yemenites** | 47 | MiddleEast | Rajeevan et al. (2003) |

**Table S1C:** Samples gathered from Europe and Asia, to be merged with the samples from Table S1B, to test hypotheses regarding Indo-European and Tibeto-Burman language dispersals into the Indian sub-continent.

**Table S2:** Shared Ancestry table between the 91 populations found in Table S1A. The matrix is ordered according to language and social group affiliations. We see that Austro-Asiatic and Tibeto-Burman populations usually show divergence from the rest of India and only cluster within themselves. Few DR_SGCs such as Paniyas, Irulas and Kadars, show little shared ancestry with other Dravidian SGC and SGBs, which is explained by their remote locations in the hills and their livelihood as nomadic hunter gatherers. The Gonds share a very high amount of ancestry with other Austro-Asiatic and Dravidian populations, which follows from linguistics, as Gondis are bilingual. The Dravidian SGB and SGAs share high ancestry with Indo-European SGA/SGB/SGCs.

| A | B | C | F3 | Err | Z |
|---|---|---|---|---|---|
| DR_SGB | IE_SGA | Gounder | -0.02328 | 0.000644 | -36.114 |
| IE_SGA | TB_SGC | Manipuri_Brahmin | -0.01583 | 0.000452 | -35.019 |
| DR_SGB | IE_SGC | Gounder | -0.02188 | 0.000657 | -33.315 |
| IE_SGA | TB_SGC | Tharu | -0.01364 | 0.000447 | -30.518 |
| DR_SGA | TB_SGC | Tharu | -0.01292 | 0.000429 | -30.084 |
| IE_SGC | TB_SGC | Tharu | -0.00843 | 0.000389 | -21.647 |
| DR_SGC | TB_SGC | Tharu | -0.00913 | 0.000436 | -20.922 |
| DR_SGC | TB_SGC | Manipuri_Brahmin | -0.0094 | 0.000484 | -19.415 |
| IE_SGA | AA_SGC | Iyer | -0.00343 | 0.000241 | -14.211 |
| IE_SGA | AA_SGC | Gond | -0.00449 | 0.000321 | -13.989 |
| DR_SGC | AA_SGC | Gond | -0.00419 | 0.000305 | -13.722 |
| IE_SGC | AA_SGC | Gond | -0.00226 | 0.000171 | -13.245 |
| IE_SGA | AA_SGC | Kol | -0.00347 | 0.000266 | -13.002 |
| IE_SGA | AA_SGC | Pallan | -0.00411 | 0.000325 | -12.638 |
| IE_SGA | AA_SGC | Bhil | -0.00326 | 0.000277 | -11.758 |
| IE_SGA | DR_SGC | Bhil | -0.0036 | 0.000343 | -10.489 |
| IE_SGC | TB_SGC | Khasi | -0.01008 | 0.001166 | -8.648 |
| IE_SGB | TB_SGC | Khasi | -0.00981 | 0.001155 | -8.49 |
| IE_SGA | AA_SGC | Chamar | -0.00292 | 0.000358 | -8.152 |
| IE_SGA | AA_SGC | Satnami | -0.00503 | 0.000853 | -5.898 |

**Table S3:** Top 10% of the significant $f_3$ statistics ($f_3$(C; A,B)) highlighting the most admixed populations in India. Gounders, Manipuri Brahmins, Tharus and Gonds are the most admixed among all tribes in India. Detailed $f_3$ statistics (for all mainland Indian populations from Table S1A available in supplementary .xlsx file SuppTable3). The Indo-European tribes such as Bhil, Kol and Chamar show signs of admixture from Austro-Asiatic tribes and Indo-European forward and SGBs. Changapa, who are a tribe in Ladakh, Jammu and Kashmir in the extreme north surprisingly show signs of admixture from Dravidian tribes, Indo-European SGAs, SGBs and tribes, showing that they have been in contact with the rest of the tribes in India. The Gondis show signs of admixture from Austro-Asiatic, Dravidian and Indo-European tribes, which is much expected as Gondis are spanned across central India. Some Gondi samples also show admixture from Indo-European and Dravidian SGAs. Expectedly, the Khasis are an admixed population from Tibeto-Burman tribes and Austro-Asiatic tribes. The Khasis are Austro-Asiatic speakers located in the northeast, along with Tibeto-Burman tribes. Notably, Manipuri Brahmins uphold the view that they are an admixed population between Indo-European and Tibeto-Burman speakers. The Tharus also are an admixed population, as had been noted earlier, with admixture from Indo-European, Dravidian, Austro-Asiatic, and Tibeto-Burman tribes, but not just Indo-Europeans.

**Table S4:** Outgroup f3 statistic results for the tests: f3(YRI; X, Y) as visualized in Figure 4 in the pie charts showing shared genetic affinity of X and Y populations, where X is an Indian population and Y is an Eurasian/southeast Asian population. The maximum f3 values are returned for every population in X w.r.t Y and then represented here in descending order. The greater the maximum shared genetic affinity, the darker is the color used in the pie chart in Fig 4. This table shows that the Europeans share more genetic drift with the IE_SGA and East Asians with the TB_SGC, reflecting on the gateways of gene flow to the Indian subcontinent.
